# An AI-assisted platform for quantitative histopathological analysis in interstitial lung disease

**DOI:** 10.64898/2026.08.16.745078

**Authors:** Ido Mizrahi, Yuanning Guo, Jiayu He, Ido Livneh, Polina Stein, Rotem Bela Shimron, Adi Raz, Maher Abu Saleh, Tina Shogan, Naama Matalon, Maya Hershfinkel, Hadas Achildiev Cohen, Ariel Shemesh, Raz Palty, Yaniv Dotan, Haguy Wolfenson, Peleg Hasson, Anas Odeh

## Abstract

Interstitial lung diseases (ILDs) are heterogeneous pulmonary disorders characterized by chronic inflammation and/or fibrosis. 30-40% of ILD patients develop fibrotic disease that is associated with progressive respiratory decline and poor prognosis, particularly in idiopathic pulmonary fibrosis. Current antifibrotic therapies slow disease progression but do not reverse fibrosis, highlighting the need for improved therapeutic strategies. Robust histopathological evaluation in preclinical models is essential for drug development; however, conventional scoring systems are semi-quantitative, labor-intensive, subject to inter-observer variability, and rely on limited field sampling. Here, we introduce FibroSight, a standalone platform for compartment-resolved quantification of lung remodeling in Sirius Red–stained sections. By integrating deep learning– based structural segmentation with color-based feature extraction, FibroSight enables highly automated whole-lobe analysis without requiring complex computational setup. The platform quantifies complementary remodeling parameters, including parenchymal collagen fraction, parenchymal tissue density, nuclear area fraction, parenchymal airspace fraction, and airway- and vascular-associated remodeling. Validated in the bleomycin-induced fibrosis model, FibroSight-derived metrics strongly correlated with expert Ashcroft scoring and showed stronger associations with histological severity than corresponding outputs from a semi-automated ImageJ-based workflow. The platform further distinguished inflammatory from fibrotic remodeling in influenza-induced lung injury and demonstrated translational proof-of-concept applicability in human ILD biopsy specimens. By enabling scalable, reproducible, and multi-compartment histological quantification, FibroSight provides a practical framework for objective assessment of lung remodeling. This approach expands conventional fibrosis evaluation by integrating fibrotic, inflammatory, airway, and vascular-associated readouts, supporting more precise analysis of disease mechanisms and therapeutic responses in preclinical and translational ILD research.

## Introduction

Interstitial lung diseases (ILDs) are a heterogeneous group of disorders characterized by inflammation and/or fibrosis, leading to progressive tissue destruction, hypoxia, and ultimately death(1,2). Collectively, ILDs contribute to an estimated 30,000 deaths annually in the United States(2). Approximately one-third of ILDs exhibit a fibrotic phenotype, with the most common form being idiopathic pulmonary fibrosis (IPF) – a chronic and rapidly progressive disease with a median survival of 3–5 years(2). Fibrotic lung diseases are driven by recurrent epithelial injury, inflammation-induced fibroblast activation, and TGF-β–mediated myofibroblast differentiation, culminating in excessive extracellular matrix (ECM) deposition and remodeling as well as progressive architectural distortion(3).

Nintedanib, pirfenidone, and nerandomilast are currently approved drugs for IPF patients(4–6), with nintedanib also approved for other forms of progressive pulmonary fibrosis (PPF) and systemic sclerosis-associated ILD(7,8). Although these agents slow the decline in lung function and appear to extend survival, they fail to halt or reverse disease progression(9–12). Both nintedanib and pirfenidone are frequently associated with gastrointestinal adverse effects and hepatotoxicity, contributing to treatment discontinuation with rates reaching up to 34% in real-world cohorts(8,13,14). Despite the recent approval of nerandomilast as a third antifibrotic agent for IPF, the development of novel therapeutic strategies to halt or reverse fibrosis remains a critical unmet need.

The gold standard for assessing the efficacy of new anti-fibrotic therapies in preclinical models is histological evaluation of lung sections(15), including the use of traditional clinical histopathological scores such as the Ashcroft scale(16,17). However, these scoring systems suffer from several inherent limitations, including substantial inter- and intra-observer variability which limits standardization across laboratories(17). Further, Ashcroft scoring is typically performed on selected microscopic fields rather than whole tissue sections and requires significant training and expertise. As a result, biologically meaningful changes in tissue architecture may be overlooked, and the comprehensive characterization of underlying pathological processes is limited.

Several computational tools have been developed to quantify fibrosis from histological lung sections aiming to address these limitations, differing substantially in methodology, automation level, accessibility, and biological resolution (Supplementary Table S1). Some approaches rely on deep learning models trained on semi-quantitative fibrosis grading scales, essentially recapitulating Ashcroft scoring. Other recent frameworks employ tile-based density indices, reporting mean tissue density and dense tissue fraction(18). However, these approaches produce single-score outputs that compress the complexity of the remodeling process, thereby limiting interpretability and sensitivity to region-specific changes(19,20). Some of the methods depend on time-consuming manual pre-processing steps, including lung tissue border detection and exclusion of blood vessels and airways, or employ semi-automated workflows. Even when integrating advanced segmentation models, these approaches typically operate on an image-by-image basis and lack batch-processing capability across large datasets, thus constraining scalability and reproducibility(21,22). In addition, several tools require proprietary software environments or complex computational setup and may be hardware-intensive, limiting accessibility and broader adoption(18,22). Collectively, these limitations highlight the need for an open-source, highly automated platform for high-resolution quantification of fibrosis that can process large-scale datasets with minimal human intervention.

Beyond parenchymal fibrosis, inflammation plays a central role in acute lung injury and ILD(23,24). However, despite the recognition of the importance of key histological features of inflammation(25), no validated, reproducible semi-quantitative scoring system with established thresholds has been developed for these parameters(26). This limitation is particularly consequential because inflammation and fibrosis are temporally overlapping and mechanistically interrelated processes that may respond differently to therapeutic intervention(27). Thus, there is major need for standardized tools that capture both inflammatory and fibrotic dimensions of injury in preclinical lung injury models.

Another underexplored aspect of preclinical lung injury assessment is vascular injury and remodeling. Pulmonary vascular alterations are increasingly recognized as both a consequence of lung damage and an active contributor to ILD progression(28–30). Distinct ILD subtypes may also exhibit different angiogenic patterns(31), and quantitative assessment of pulmonary vascular volume on CT imaging has been identified as a strong predictor of mortality in IPF(32). Despite this, vascular parameters are seldom incorporated into routine preclinical histological assessment. While three-dimensional imaging modalities can provide detailed vascular characterization, they are not readily scalable to routine preclinical workflows(33,34). Standard histological sections, however, remain widely available and can yield meaningful two-dimensional vascular and perivascular morphometric readouts when analyzed systematically. Consequently, there is an opportunity to extract vascular-associated remodeling features from routine histological preparations.

To address all the above, we introduce FibroSight, a standalone quantitative platform that enables streamlined, compartment-resolved assessment of lung remodeling from histological images. FibroSight integrates automated structural segmentation with color-based feature extraction to generate complementary metrics describing parenchymal collagen fraction, tissue density, nuclear area fraction, and both vascular and airway remodeling from Sirius Red–stained lung sections. Together, these parameters provide an integrated quantitative representation of structural and cellular remodeling processes that are difficult to capture comprehensively using conventional manual or semi-quantitative approaches in ILDs and other diseases leading to lung injury.

## Results

To enable quantitative assessment of lung remodeling across entire histological sections, FibroSight was designed as a hierarchical workflow that first defines lung tissue boundaries and then extracts compartment-specific remodeling features. By integrating deep learning–based structural detection with color space–driven feature extraction, FibroSight partitions the lung into distinct anatomical compartments, specifically isolating the parenchyma from blood vessels and bronchi. Sirius Red staining is well suited for this workflow because it binds strongly to collagen fibers, is simple and reproducible, and can be readily quantified by image analysis. Together, these steps generate a set of biologically interpretable metrics, including parenchymal collagen fraction, parenchymal tissue density, and nuclear area fraction, which are organized into standardized primary measurements and derived parameters (Supplementary Table S2). These outputs form the foundation for the comparative analyses and validation steps presented throughout this study (Fig. 1).

**Figure 1.**
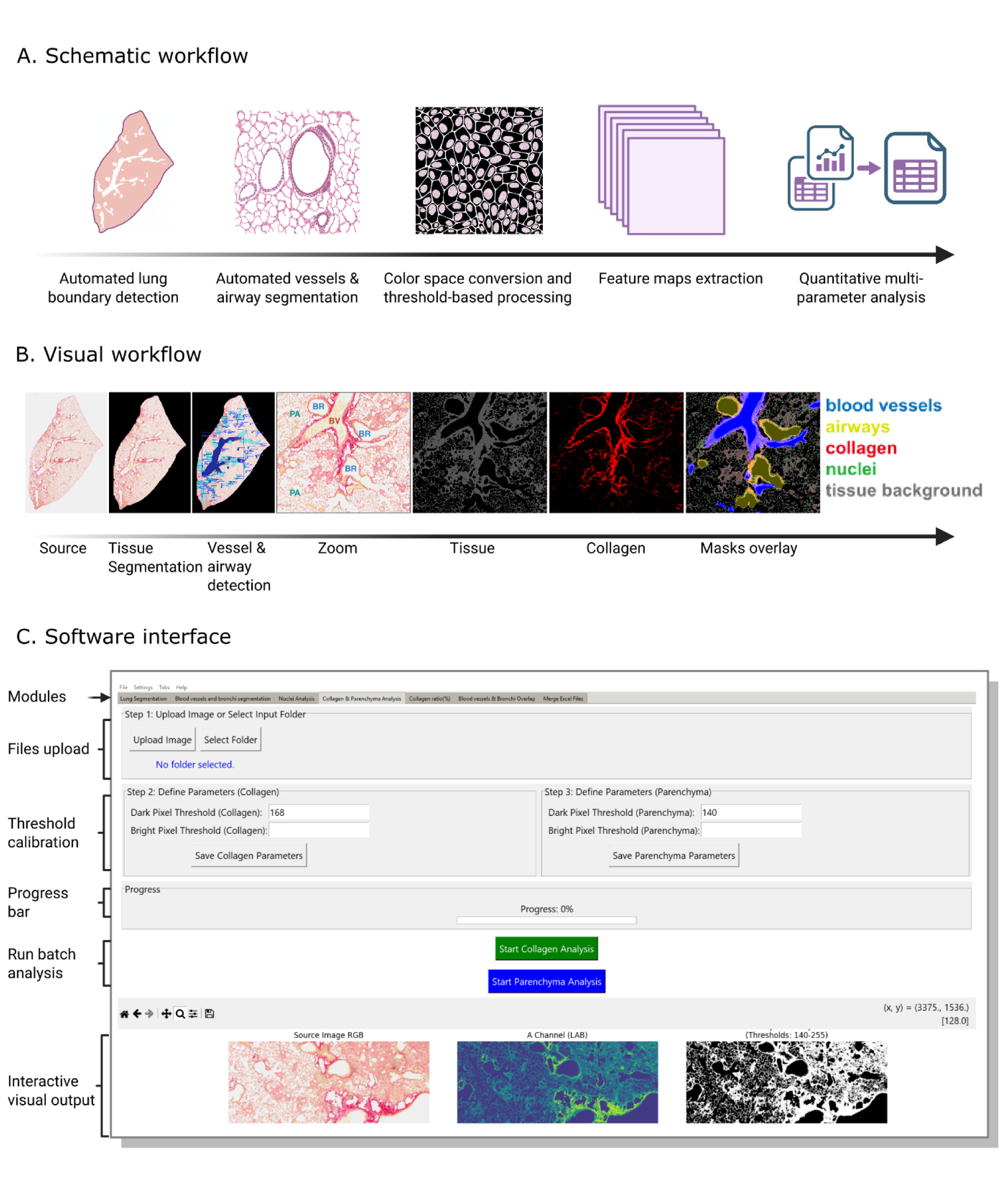
Overview of the FibroSight workflow and extracted histopathological features. **(A)** Schematic workflow. FibroSight performs highly automated analysis of histological sections through a sequential pipeline that includes YOLOv11-based lung border detection and automated segmentation of blood vessels and airways, followed by color-space conversions with threshold-based processing, and extraction of high-resolution feature maps. These steps enable scalable quantification of key histological parameters, including tissue density, fibrosis, and cellularity across entire lung sections. The schematic illustration of lung border detection in panel A was generated with assistance from Google Gemini. **(B)** Representative visual workflow. Example outputs illustrating the main processing stages. Starting from the original Sirius Red–stained lung section, FibroSight performs automated border segmentation as well as detection of vessels and airways, followed by color-based segmentation of tissue, collagen and nuclei masks. The resulting segmentation masks are combined to generate compartment-resolved maps of lung tissue, vessels, airways, collagen, and nuclei, enabling extraction of multiple quantitative metrics describing structural and cellular remodeling within the lung. **(C)** The software’s interface. The analytical pipeline consists of 7 modules accessible via the tab bar. Each module analysis begins with uploading files: first, individual images for threshold calibration, and then the entire input folder. Next, the researcher can interactively define a threshold and examine its accuracy by looking at the visual output panel, which instantly updates whenever a new threshold is set.

### FibroSight enables compartment-specific detection of fibrotic remodeling in bleomycin-treated lungs

To evaluate the platform’s performance in quantifying fibrosis-associated lung tissue remodeling, FibroSight was applied to analyze lungs from saline- and bleomycin-treated mice, a well-established experimental model for lung injury and fibrosis(35,36). Representative Sirius Red– stained sections demonstrated characteristic bleomycin-induced remodeling, including marked tissue consolidation, reduced alveolar airspace, architectural distortion, focal parenchymal collagen accumulation, and cell-dense remodeled regions (Fig. 2A–F).

**Figure 2.**
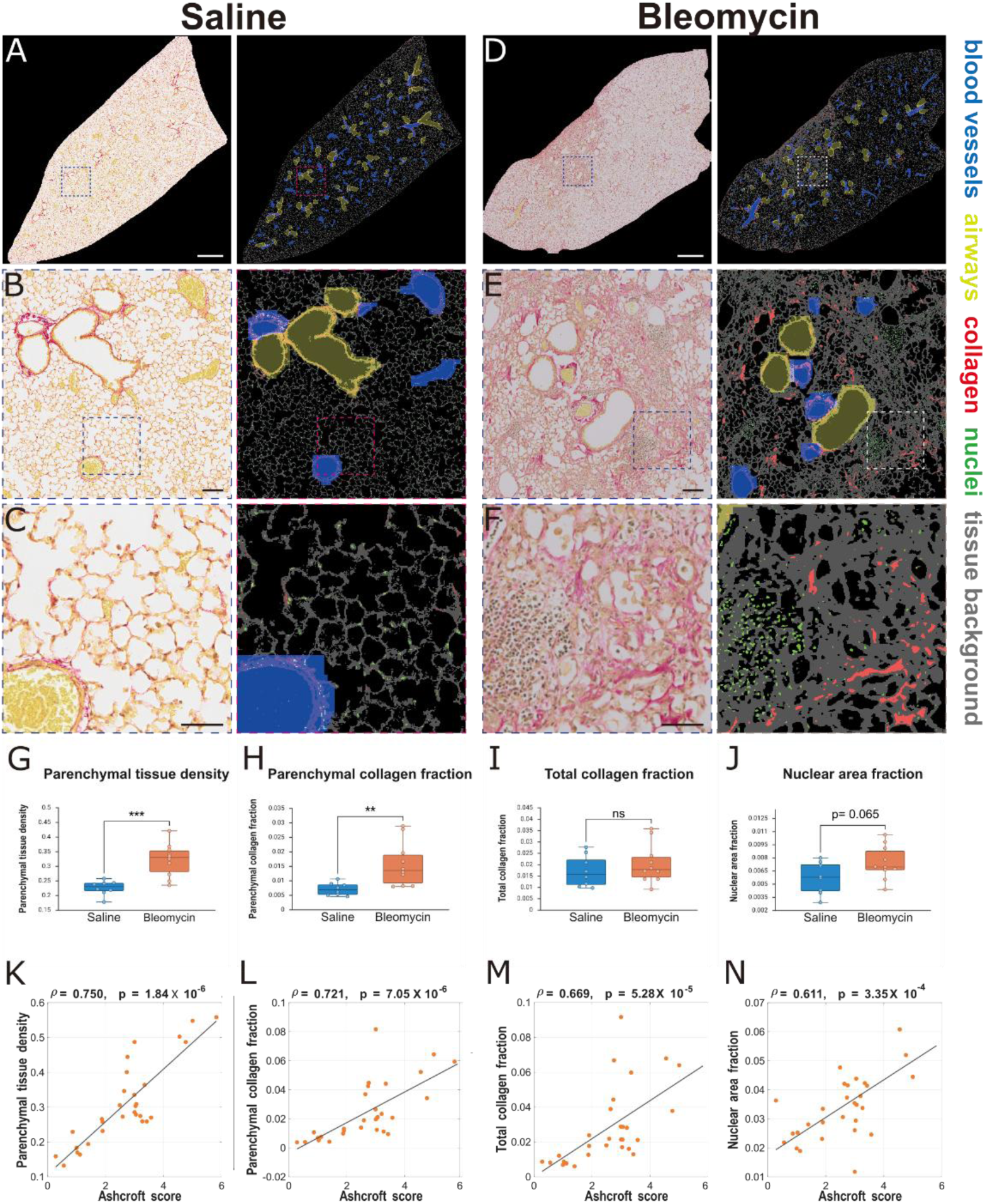
FibroSight quantifies bleomycin-induced lung remodeling and aligns with expert histopathological scoring. **(A–F)** Representative Sirius Red–stained lung lobe images from saline-treated (A–C) and bleomycin-treated (D–F) mice with corresponding FibroSight mask overlays. In each pair, the original histological image is shown on the left and the FibroSight visual output is shown on the right. Gray indicates tissue/background, red collagen, green nuclei, blue blood vessels, and yellow airways. Lower panels show magnified views of the boxed regions. Bleomycin-treated lungs demonstrate increased tissue density, parenchymal collagen deposition, architectural distortion, and focal cellular infiltrates compared with saline-treated controls. Scale bars: 1 mm in whole-lobe images and 100 μm in magnified images. **(G–J)** Quantitative comparison of FibroSight-derived parameters between saline and bleomycin-treated mice. Bleomycin injury significantly increased parenchymal tissue density (G) and parenchymal collagen fraction (H), indicating loss of airspace and collagen accumulation within the gas-exchange compartment. In contrast, total collagen fraction (I) did not differ significantly between groups, highlighting the lower sensitivity of global collagen measurements to spatially heterogeneous fibrosis. Nuclear area fraction (J) showed a trend toward increased cellularity following bleomycin injury but did not reach statistical significance. Each point represents one mouse, with values calculated as area-weighted averages across all analysed lung lobe images from the same animal. Statistical significance was assessed using an unpaired t-test. **(K–N)** Image-level correlations between expert pathologist-assigned Ashcroft scores and FibroSight-derived quantitative parameters across lung lobe images from the bleomycin-induced lung injury model. Parenchymal tissue density showed the strongest correlation with Ashcroft score (ρ = 0.750, p = 1.84 × 10⁻⁶; K), followed by parenchymal collagen fraction (ρ = 0.721, p = 7.05 × 10⁻⁶; L), total collagen fraction (ρ = 0.669, p = 5.28 × 10⁻⁵; M), and nuclear area fraction (ρ = 0.611, p = 3.35 × 10⁻⁴; N). Each point represents one lung lobe image. Solid lines indicate linear trend lines shown for visualization only; correlation coefficients and p values were calculated using Spearman’s rank correlation.

Quantitative analysis revealed a significant increase in parenchymal tissue density in bleomycin-treated lungs relative to saline controls (Fig. 2G), indicating consolidation of the alveolar airspaces following fibroproliferative processes. Parenchymal collagen fraction, defined as the area of collagen deposition within the alveolar parenchyma normalized to clean lung area (net lung area excluding major blood vessels and bronchi), was significantly elevated following bleomycin injury (Fig. 2H). In contrast, the total collagen fraction, calculated as total collagen area normalized to lung area, did not significantly differ between groups (Fig. 2I). This discrepancy suggests that global collagen measurements may be less sensitive to spatially heterogeneous parenchymal fibrosis, particularly when collagen-rich non-parenchymal structures such as large vessels and bronchi are included in the analyzed area.

We also assessed the nuclear area fraction, calculated as total nuclear area normalized to lung area, as a marker of overall tissue cellularity. Bleomycin-treated lungs showed a modest increase in nuclear area fraction. Still, this trend did not reach statistical significance (Fig. 2J). This is consistent with the time point analyzed, in which bleomycin injury is expected to be dominated by fibroproliferative remodeling rather than peak inflammatory infiltration. Together, these findings show that FibroSight-derived parenchyma-specific metrics capture bleomycin-induced structural remodeling more sensitively than global collagen or cellularity-based measurements.

### Validation against expert histopathological scoring

To validate FibroSight against expert histopathological assessment, we compared its automated outputs with Ashcroft scores assigned by two blinded, independent lung pathologists on 30 Sirius Red–stained lung lobe images from the bleomycin model. Spearman rank correlation analysis was employed to assess the relationship between FibroSight-derived parameters and Ashcroft scores.

Among the evaluated metrics, parenchymal tissue density showed the strongest correlation with Ashcroft scoring (ρ = 0.75, p < 0.001, Fig. 2K). Parenchymal collagen fraction also correlated strongly with Ashcroft score (ρ = 0.721, p < 0.001, Fig. 2L) while total collagen fraction showed a significant but weaker association (ρ = 0.669, p < 0.001, Fig. 2M).

Nuclear area fraction, a cellularity-related parameter not directly addressed by the Ashcroft score, also correlated with pathologist assessment (ρ = 0.611, p < 0.001, Fig. 2N). These results indicate that FibroSight captures key quantitative features associated with expert-defined fibrosis severity while providing additional continuous, compartment-resolved readouts beyond a single ordinal score.

In contrast, vascular and bronchial wall area fractions (Fig. S1) exhibited moderate correlations with Ashcroft scores. This is consistent with the parenchyma-focused nature of Ashcroft scoring, which excludes fields predominantly occupied by large bronchi or blood vessels and grades fibrosis only within the remaining gas-exchange compartment(16).

### Benchmarking FibroSight against a semi-automated ImageJ workflow

To benchmark FibroSight against an established quantitative histopathology approach, we compared its outputs with those obtained using a recently described semi-automated ImageJ-based fibrosis quantification plugin(21). This workflow provides quantitative measurements of fibrosis and airspace in lung histology and was therefore used as a comparator for shared FibroSight-derived parameters. Benchmarking was performed on the image set used for expert histopathological validation, comprising 30 Sirius Red–stained lung sections from the bleomycin model and spanning a broad range of fibrosis severities. Each image represented a single lung lobe, enabling direct comparison between computational outputs and expert Ashcroft scoring.

Direct comparison of shared parameters revealed strong agreement between FibroSight and the ImageJ plugin. FibroSight-derived parenchymal collagen fraction strongly correlated with ImageJ-derived fibrosis measurements (ρ = 0.879, p < 0.001; Fig. 3A), and FibroSight-derived parenchymal airspace fraction, the inverse measure of parenchymal tissue density, strongly correlated with ImageJ-derived airspace measurements (ρ = 0.958, p < 0.001; Fig. 3B). These findings indicate that FibroSight captures key structural features of lung remodeling, in agreement with an established semi-automated analysis workflow.

**Figure 3.**
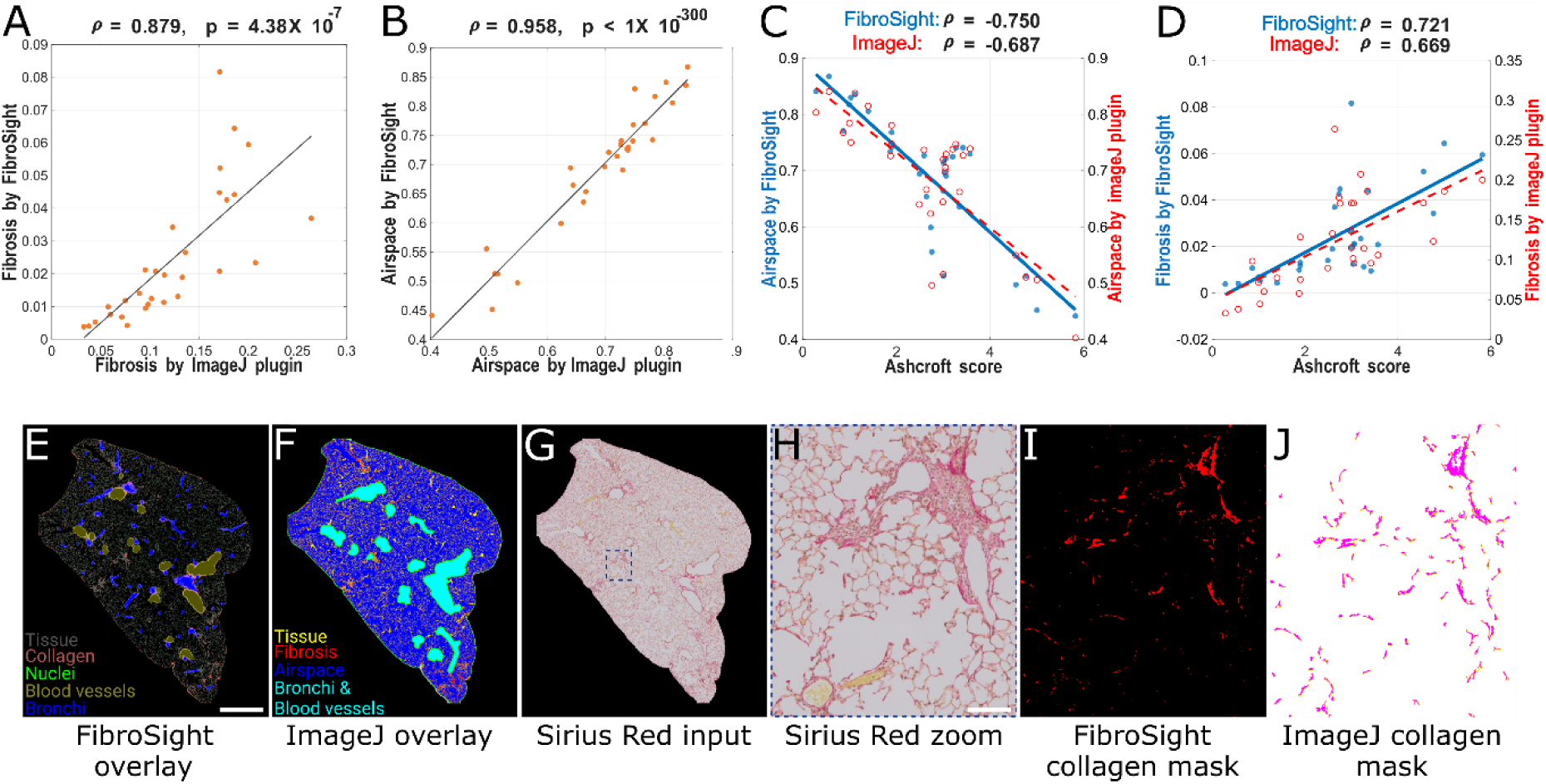
Benchmarking FibroSight against a semi-automated ImageJ-based fibrosis quantification workflow. **(A, B)** Correlation between shared quantitative outputs generated by FibroSight and the ImageJ plugin across a matched subset of 30 Sirius Red–stained lung lobe images. FibroSight-derived parenchymal collagen fraction strongly correlated with ImageJ-derived fibrosis measurements (A), and FibroSight-derived parenchymal airspace fraction strongly correlated with ImageJ-derived airspace measurements (B). **(C, D)** Correlation of FibroSight- and ImageJ-derived parameters with expert Ashcroft scores in the same image subset. Parenchymal airspace fraction showed a stronger inverse association with Ashcroft score when quantified by FibroSight than by the ImageJ plugin (C). Similarly, FibroSight-derived parenchymal collagen fraction showed a stronger association with Ashcroft score than ImageJ-derived fibrosis measurements (D). Blue indicates FibroSight-derived metrics and red indicates ImageJ-derived metrics. Each image represents a single lung lobe, and the same 30 images were used for expert pathologist validation. **(E–J)** Representative visual comparison of FibroSight and ImageJ plugin outputs. FibroSight composite mask overlay showing tissue, collagen, nuclei, blood vessels, and airways (E) scale bar: 1 mm. Representative ImageJ plugin output following manual preprocessing, including manual curation of the tissue border and manual annotation of airspace/ductal regions and blood vessels, with fibrosis-positive areas shown in the resulting output (F). The corresponding Sirius Red input image (G), magnified region of interest (H), FibroSight-derived collagen mask (I), and ImageJ-derived collagen/fibrosis mask (J) are shown. Scale bar: 100 μm.

We next evaluated alignment with expert histopathological assessment. While both tools showed significant correlations with Ashcroft scores, FibroSight-derived airspace and fibrosis metrics exhibited stronger correlations with expert scoring than the corresponding ImageJ-derived measurements (Fig. 3C, D). Representative outputs demonstrated broadly similar segmentation patterns between the two workflows (Fig. 3E–J). However, the workflows differed substantially in the level of manual intervention required. The ImageJ-based workflow required image-by-image manual preprocessing, including curation of tissue borders and manual annotation of airspace/ductal regions and blood vessels, before downstream quantification. In contrast, FibroSight performed automated tissue, airway, and vessel segmentation and enabled batch analysis following the initial threshold calibration step. Together, these findings indicate that FibroSight preserves agreement with established quantitative outputs while reducing image-by-image manual processing.

### FibroSight reveals size-dependent pulmonary vascular remodeling in bleomycin-induced lung injury

Given that FibroSight inherently quantifies vascular morphometric readouts, we investigated its potential to detect vascular remodeling associated with fibrotic lung injury. To that end, we quantified blood vessel morphometry in Sirius Red–stained lung sections from saline- and bleomycin-treated mice. Automated vessel segmentation facilitated the high-throughput extraction of vessel-level features, including equivalent diameter, cross-sectional area, axis lengths, eccentricity, and other geometric descriptors (Supplementary Table S3).

Representative whole-lobe images and higher-magnification regions, together with corresponding FibroSight tissue and vessel masks, demonstrated reliable identification of vascular profiles within the lung tissue compartment (Fig. 4A–D). In bleomycin-treated lungs, these outputs illustrated altered vascular distribution across remodeled tissue regions. Analysis of vessel equivalent diameter revealed a marked shift in the vascular size distribution in bleomycin-treated lungs compared with saline controls (Kolmogorov–Smirnov test, p = 3.4 × 10⁻⁷⁰; Fig. 4E). This shift was characterized by a relative increase in smaller vessel profiles and a reduction in mid-sized and larger vascular structures.

**Figure 4.**
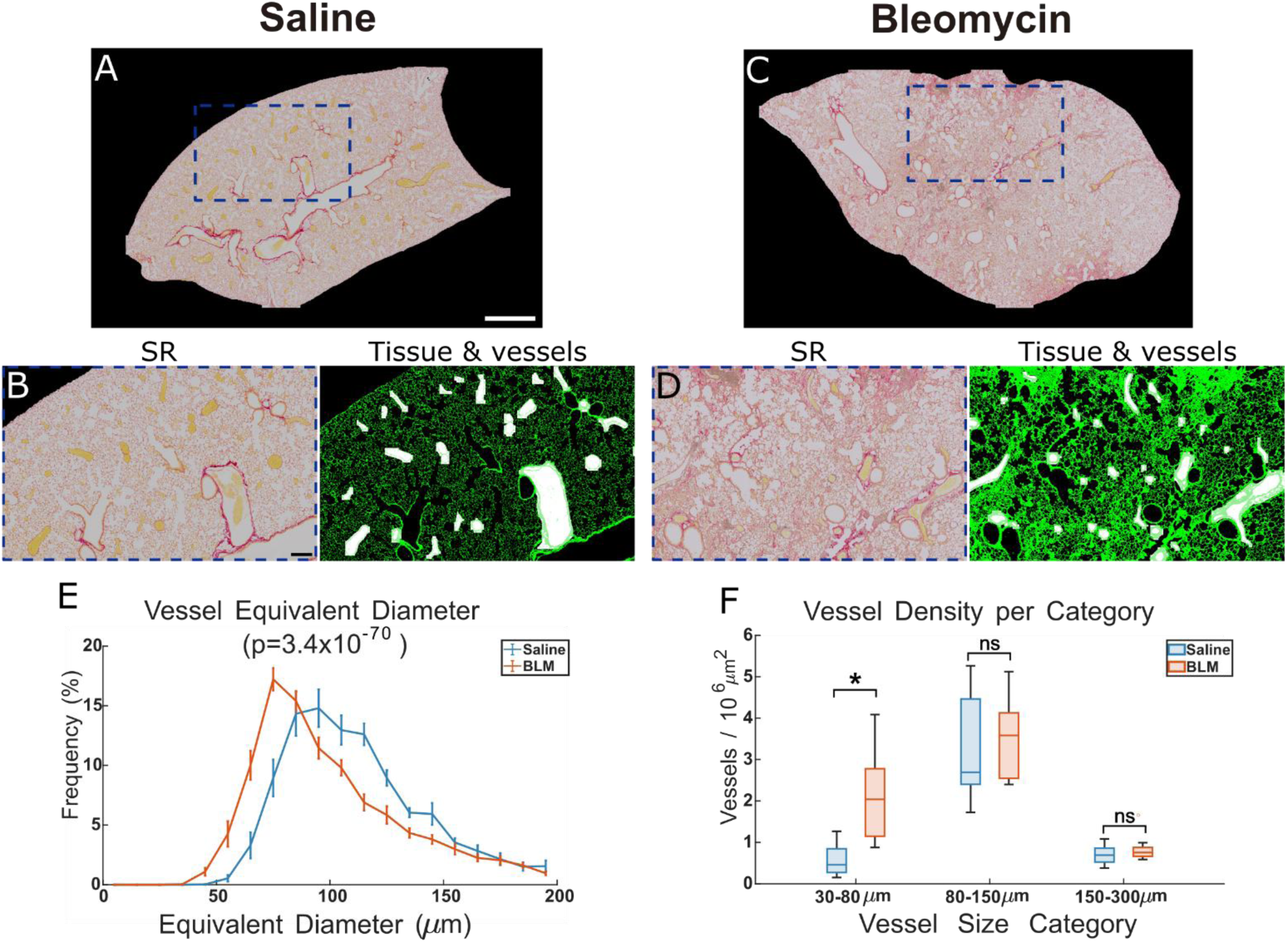
FibroSight detects size-dependent vascular remodeling in the bleomycin lung injury model. **(A–D)** Representative Sirius Red–stained lung sections from saline-treated (A,B) and bleomycin-treated (C,D) mice. Whole-lobe images are shown in (A,C), with dashed blue boxes indicating the regions shown at higher magnification in (B,D). Magnified regions are shown as Sirius Red input images together with corresponding FibroSight visual outputs, in which tissue masks are shown in green and blood vessel masks in white. Scale bars: 1mm in whole lobe images and 100 µm in magnified images. **(E)** Frequency distribution of segmented vessel equivalent diameters below 200 µm. Bleomycin injury induced a significant shift in vessel profile size distribution, with increased relative frequency of smaller vessel profiles compared with saline controls (Kolmogorov–Smirnov test, p = 3.4 × 10⁻⁷⁰). **(F)** Vessel profile density stratified by equivalent diameter category and normalized to parenchymal area (vessels/10⁶ µm²). Bleomycin-treated lungs showed increased vessel density in the 30–80 µm diameter range. Statistical comparisons were performed between saline and bleomycin groups within each size category.

To determine whether this shift in size distribution was accompanied by an absolute increase in vessel abundance, we stratified segmented vessel profiles into discrete diameter classes and normalized their number to parenchymal area. This analysis revealed a significant increase in vessel density specifically within the 30–80 µm diameter range in bleomycin-treated lungs, whereas densities of larger vessel profiles, including the 80–150 µm and 150–300 µm ranges, did not differ (Fig. 4F). These findings are in line with the heterogeneous vascular remodeling described in IPF, including reports of altered angiogenic patterns, expansion of small vascular structures, and vascular remodeling within fibrotic regions(31,37).

### FibroSight distinguishes inflammatory from fibrotic lung remodeling in influenza-induced tissue damage

To evaluate whether FibroSight can capture histopathological remodeling associated with other inflammatory lung injuries, we applied the platform to Sirius Red–stained lung sections from influenza-infected mice. Representative lung images and higher-magnification views illustrate diffuse architectural disruption, marked cellular infiltration, increased tissue density, and airway-associated remodeling in influenza-infected lungs compared with preserved alveolar structure in saline-treated controls (Fig. 5A–F). High-magnification views further demonstrated leukocyte-rich intra-alveolar and interstitial inflammatory infiltrates within affected regions.

**Figure 5.**
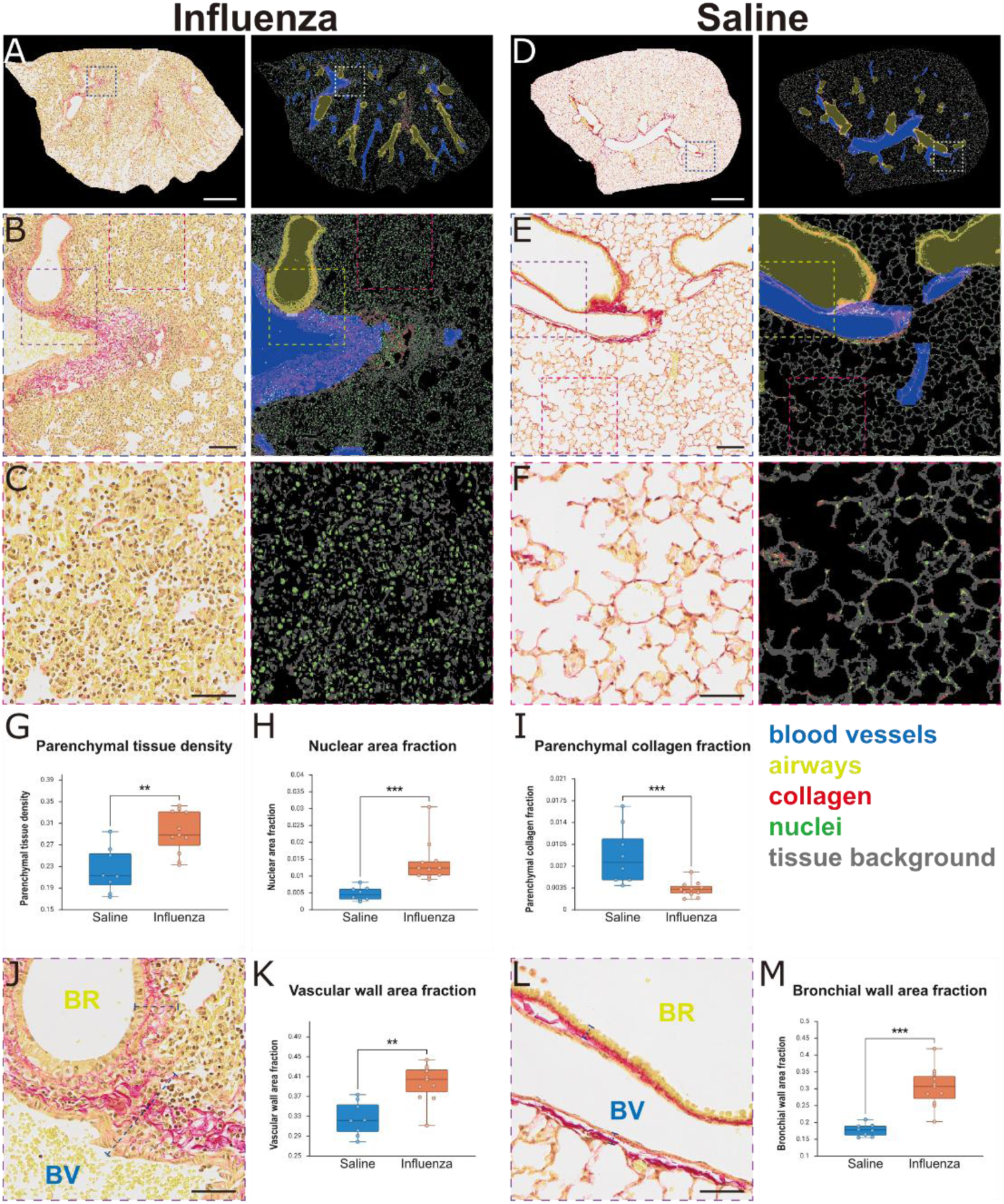
FibroSight identifies inflammation-dominant remodeling in influenza-induced tissue damage. **(A–F)** Representative Sirius Red–stained lung sections and corresponding FibroSight mask overlays from influenza-infected (A–C) and saline-treated control mice (D–F). Whole-lobe images are shown in (A, D), airway/vascular regions in (B, E), and higher-magnification parenchymal regions in (C, F). Mask overlays indicate blood vessels in blue, airways in yellow, collagen in red, nuclei in green, and tissue/background in gray. Influenza-infected lungs show diffuse cellular infiltration, increased tissue density, disruption of alveolar architecture, and airway- and vessel-associated remodeling compared with saline controls. Scale bars: 1 mm in whole-lobe images and 100 μm in magnified images. **(G–I)** FibroSight quantification of parenchymal remodeling. Influenza infection increased parenchymal tissue density (G) and nuclear area fraction (I), while reducing parenchymal collagen fraction (H), consistent with inflammation-dominant tissue injury involving disruption of collagen-containing alveolar septa rather than collagen-dominant fibrosis. **(J–M)** Representative airway and vascular regions from influenza-infected (J) and saline-treated mice (L), with corresponding quantification of vascular wall area fraction (K) and bronchial wall area fraction (M). Influenza infection increased both vascular and bronchial wall area fractions, indicating airway- and vessel-associated inflammatory remodeling. Each point represents an individual mouse, with lobe-level measurements averaged and weighted by relative lobe area. Statistical comparisons were performed using Mann–Whitney tests.

Quantitative analysis revealed a significant increase in parenchymal tissue density in influenza-infected lungs relative to saline-treated controls (Fig. 5G), reflecting widespread inflammatory consolidation and loss of alveolar airspace. In parallel, nuclear area fraction was markedly elevated in the influenza-infected lungs (Fig. 5H), indicating increased tissue cellularity consistent with inflammatory infiltration. Together, these metrics describe a predominant cellular and inflammatory remodeling phenotype, distinct from the collagen-driven remodeling observed in the bleomycin model.

In contrast to bleomycin-induced fibrosis, parenchymal collagen fraction was significantly reduced in influenza-infected lungs (Fig. 5I). This finding aligns with the disruption or loss of alveolar septal collagen and the degradation of parenchymal architecture rather than collagen accumulation. Such matrix disruption is a recognized feature of the peak inflammatory phase of influenza-induced inflammation and has been linked to increased extracellular matrix proteolysis mediated by inflammatory cell–derived matrix metalloproteinases (MMPs)(38–41). Consistent with this mechanism, we observed massive inflammatory infiltration, a potential source of collagen-degrading MMPs within the alveolar spaces and parenchymal tissue of infected lungs (Fig. 5C, F).

Beyond parenchymal remodeling, FibroSight detected significant increases in bronchial wall area fraction and vascular wall area fraction in influenza-infected lungs compared with saline controls (Fig. 5J–M). These changes likely reflect airway- and vessel-associated inflammatory remodeling, including wall-associated cellular infiltration, edema, and structural thickening.

Because FibroSight-derived wall fractions are based on two-dimensional histological segmentation, they do not distinguish specific anatomical wall layers, such as intima, media, or adventitia. Nevertheless, these parameters provide quantitative compartment-specific readouts of airway and vascular involvement in inflammatory lung injury.

To further evaluate whether FibroSight-derived parameters reflect graded inflammation severity, we compared lobe-level FibroSight outputs with an adapted semi-quantitative inflammatory scoring system based on alveolar/interstitial, perivascular, and peribronchial inflammatory involvement. Several FibroSight parameters showed significant associations with inflammatory score (Supplementary Fig. S2A). Nuclear area fraction showed the strongest positive correlation with inflammation severity (ρ = 0.661, p < 0.001), consistent with increasing cellular infiltration. Parenchymal tissue density also correlated positively with inflammatory score (ρ = 0.636, p < 0.001), reflecting progressive tissue consolidation during inflammatory injury. Bronchial wall area fraction was positively associated with inflammatory score (ρ = 0.534, p < 0.001), supporting airway-associated inflammatory remodeling. Collagen-related parameters showed weaker positive correlations, while vascular wall area fraction was not significantly associated with inflammation severity.

Severity-stratified analysis further demonstrated quantitative differentiation between inflammation groups for several parameters, with severe inflammation showing increased nuclear area fraction, parenchymal tissue density, and bronchial wall area fraction compared with lower-severity groups (Supplementary Fig. S2B). Together, these findings indicate that FibroSight can capture graded histological inflammation severity, with nuclear area fraction and parenchymal tissue density emerging as the most informative quantitative readouts in the influenza model.

Collectively, these results demonstrate that FibroSight resolves distinct remodeling signatures across lung injury contexts, distinguishing cellular-dominant inflammatory remodeling in influenza infection from collagen-dominant fibrotic remodeling in the bleomycin model.

### FibroSight demonstrates proof-of-concept applicability in human ILD biopsy specimens

To assess the translational potential of FibroSight, we applied the platform to human ILD biopsy specimens, including transbronchial biopsies (TBB) and surgical wedge-resection specimens. These specimen types present distinct analytical challenges. TBB specimens are smaller and frequently affected by tissue compression, fragmentation, and architectural distortion, which complicates spatial segmentation of intrapulmonary airways and vasculature. In contrast, wedge-resection specimens provide larger, architecture-preserved tissue sections that more closely resemble the whole-lobe sections analyzed in preclinical models, allowing more comprehensive assessment of compartment-resolved segmentation in human tissue.

We first evaluated FibroSight in human transbronchial biopsy specimens, representing a clinically relevant but technically challenging tissue format. This cohort included 16 patients, represented by 16 paraffin blocks, from which 48 cropped tissue-region images were generated for analysis. FibroSight reliably delineated tissue boundaries within the cropped regions and enabled extraction of tissue, collagen, and nuclear features. However, automated blood vessel and bronchus segmentation was unreliable in this specimen type, consistent with reduced tissue volume and loss of preserved anatomical landmarks. Accordingly, downstream TBB analysis focused on tissue-level and color-based parameters rather than compartment-resolved vascular or airway measurements (Fig. S3). These findings support the feasibility of FibroSight-based quantitative feature extraction from limited biopsy material, while identifying structural segmentation as a key area for TBB-specific adaptation.

We next evaluated FibroSight in surgical wedge-resection specimens, where preserved tissue architecture enabled assessment of both color-based feature extraction and spatial compartment segmentation (Fig. 6). These samples were obtained from three patients and represented 12 paraffin blocks in total. Because wedge specimens contained large and heterogeneous tissue areas, each digitized slide was divided into anatomically coherent, non-overlapping tissue-region images comparable in scale to murine lung lobe images, yielding 32 image-level regions for analysis. Each region was assigned an independent Ashcroft-based fibrosis score and analyzed using FibroSight.

**Figure 6.**
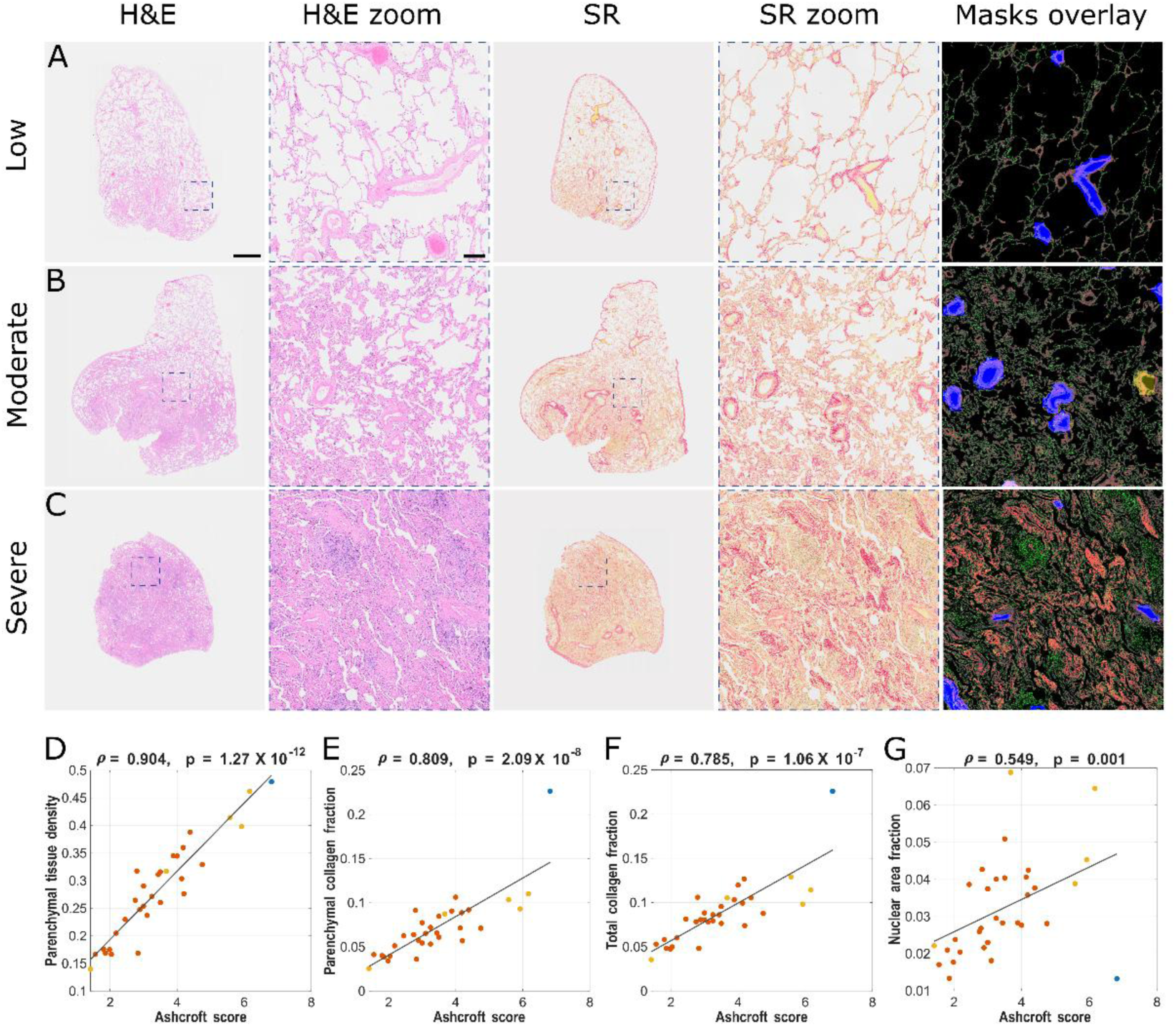
FibroSight quantifies histological remodeling severity in human surgical wedge biopsy specimens. **(A–C)** Representative human wedge-resection tissue-region images stratified by Ashcroft-based fibrosis severity: low (A), moderate (B), and severe (C). For each severity category, H&E-stained images, corresponding H&E magnified regions, Sirius Red (SR)-stained images, SR magnified regions, and FibroSight-derived mask overlays are shown. In the mask overlays, gray indicates tissue/background, red collagen, green nuclei, blue blood vessels, and yellow airways. Representative regions illustrate progressive architectural distortion, increased tissue density, and collagen-rich remodeling with increasing histological severity. Scale bars: 1 mm in whole tissue-region images and 200 μm in magnified images. **(D–G)** Image-level correlations between FibroSight-derived quantitative parameters and expert Ashcroft-based fibrosis scores across wedge-resection tissue-region images. Parenchymal tissue density showed the strongest association with histological severity (ρ = 0.904, p = 1.27 × 10⁻¹²; D), followed by parenchymal collagen fraction (ρ = 0.809, p = 2.09 × 10⁻⁸; E) and total collagen fraction (ρ = 0.785, p = 1.06 × 10⁻⁷; F). Nuclear area fraction showed a more moderate correlation with Ashcroft score (ρ = 0.549, p = 0.001; G). Each point represents one tissue-region image derived from 12 paraffin blocks from three patients; point colors indicate patient identity. Correlations were calculated using Spearman’s rank correlation, and gray lines indicate linear trend lines shown for visualization only.

Representative tissue-region images spanning low, moderate, and severe Ashcroft-based fibrosis severity are shown in Fig. 6A–C. H&E and Sirius Red images demonstrated progressive architectural distortion, increased tissue density, and collagen-rich remodeling with increasing fibrosis severity. FibroSight delineated tissue boundaries across wedge-derived images and identified blood vessels and bronchi in most specimens. Blood vessel segmentation was more consistent than bronchial segmentation; however, minor automated misidentifications were observed. To ensure high-fidelity downstream quantification, automated masks were reviewed and manually curated when necessary, including addition of missed microvasculature and removal of falsely segmented bronchial structures (Supplementary Table S4).

Following curation, FibroSight-derived parenchymal tissue density showed the strongest image-level correlation with expert Ashcroft-based fibrosis scores (ρ = 0.904, p < 0.001; Fig. 6D).

Parenchymal collagen fraction also correlated strongly with fibrosis severity (ρ = 0.809, p < 0.001; Fig. 6E), whereas total collagen fraction showed a slightly weaker but still robust association with Ashcroft scores (ρ = 0.785, p < 0.001; Fig. 6F), reinforcing the value of compartment-resolved analysis. Nuclear area fraction showed a more moderate correlation with Ashcroft score (ρ = 0.549, p = 0.001; Fig. 6G), likely reflecting the more complex and context-dependent relationship between cellularity, inflammation, and fibrosis in human ILD tissue.

Together, these findings demonstrate that FibroSight captures key quantitative features of local histological remodeling severity in architecture-preserved human biopsy specimens, supporting its translational applicability to human ILD tissue.

## Discussion

Here, we introduce FibroSight, a platform developed to streamline histological analysis workflows in ILD. By integrating deep learning–based structural segmentation with color-based feature extraction, FibroSight provides an automated, compartment-resolved framework for quantitative analysis of Sirius Red-stained lung sections. The platform quantifies complementary parameters, including fibrosis, parenchymal density, cellularity, and airway and vascular remodeling, capturing distinct yet interconnected pathological processes relevant across a broad spectrum of ILDs and preclinical lung injury models(42–44). We validated FibroSight against expert histopathological assessment, benchmarked it against a semi-automated ImageJ workflow, and demonstrated its applicability across distinct preclinical disease models and human lung biopsies.

Sirius Red staining was selected as the histological basis for FibroSight because it is widely used for fibrosis evaluation, binds strongly to collagen fibers, and provides a simple, reproducible staining workflow compatible with quantitative image analysis. Compared with Masson’s trichrome, which may show variable staining intensity, and collagen immunohistochemistry, which provides molecular specificity but requires antibody-based optimization, Sirius Red offers a practical balance between collagen visualization, reproducibility, and scalability. A detailed Picrosirius Red staining protocol is provided in the Supplementary Methods.

Benchmarking FibroSight against a semi-automated ImageJ-based fibrosis workflow demonstrated high concordance between shared parameters. Notably, FibroSight showed stronger correlations with Ashcroft scoring for corresponding fibrosis-related metrics, while substantially reducing manual preprocessing steps, including tissue border curation and exclusion of vessels and bronchi. Although threshold calibration remains user-defined, it is performed once per dataset and preserves human supervision while helping accommodate staining variability. Following this initial calibration, FibroSight enables batch analysis with minimal image-by-image intervention, improving scalability and reproducibility across large cohorts. A related advantage is that FibroSight analyzes entire lung lobe images rather than selected fields of view used in semi-quantitative Ashcroft scoring. This whole-image approach minimizes field-selection bias, improves representation of heterogeneous disease regions, and, when combined with compartment-resolved analysis, enables more standardized quantification of disease severity across experimental groups and treatments.

In the bleomycin-induced fibrosis model, FibroSight revealed a marked increase in parenchymal collagen fraction, whereas total collagen fraction was minimally affected. This divergence likely reflects the dilution of disease-associated parenchymal collagen by non-parenchymal collagen-rich structures such as bronchi and large vessels when using global lung measurements. Automated segmentation of bronchi and blood vessels enabled fibrosis assessment focused specifically on the gas-exchange compartment. Bleomycin-treated lungs also exhibited increased parenchymal tissue density, with this parameter showing the strongest association with expert Ashcroft scoring. Parenchymal collagen fraction also correlated strongly with Ashcroft scores, and together, these parameters capture complementary aspects of fibrotic remodeling. In contrast, nuclear area fraction did not differ significantly between bleomycin-treated and control lungs, suggesting that at the administered dose and evaluated time point, remodeling was predominantly matrix-driven rather than inflammation-driven. Nevertheless, nuclear area fraction correlated with Ashcroft scores, despite cellular burden not being explicitly quantified within the Ashcroft framework. This suggests that cellularity may still contribute to overall histopathological severity and highlights the added value of measuring cellular and fibrotic features simultaneously.

FibroSight also enabled quantitative characterization of vascular remodeling. In bleomycin-treated lungs, we detected a significant shift in the distribution of blood vessel equivalent diameters, reflecting remodeling of the vascular tree at the level of individual vessels. Such alterations suggest active remodeling of the pulmonary microvasculature during fibrotic injury, a process increasingly recognized as a key component of ILD pathogenesis. These findings are consistent with growing evidence that vascular alterations are integral to fibrotic lung disease(31,45–47). Although 2D histology cannot fully represent the 3D complexity of the pulmonary vasculature, FibroSight provides a practical and accessible approach for systematic vascular quantification across large image datasets. Future studies using advanced 3D techniques will be important for further validating and expanding these observations.

Furthermore, analysis of the influenza model revealed a remodeling phenotype distinct from bleomycin-induced fibrosis. Influenza-infected lungs demonstrated increased parenchymal tissue density and elevated nuclear area fraction, consistent with inflammatory consolidation and immune cell infiltration, while parenchymal collagen fraction was reduced (in contrast to the bleomycin model), likely reflecting alveolar septal disruption during acute inflammation. These findings demonstrate FibroSight’s capacity to distinguish cellular-dominant inflammatory injury from collagen-dominant fibrotic remodeling. Further, FibroSight also enabled assessment of airway and vascular structural changes in the inflammatory setting, showing increased wall-associated remodeling in influenza-infected lungs. These changes likely reflect airway- and vessel-associated inflammatory infiltration, edema, and structural thickening. Further studies incorporating compartment-specific markers are needed to define the affected structural layers and underlying mechanisms.

Finally, to explore translational applicability, we applied FibroSight to human ILD biopsy specimens. Performance was strongest in surgical wedge specimens, where preserved tissue architecture enabled extraction of tissue, collagen, nuclear, airway, and vascular features, and FibroSight-derived parenchymal tissue density and collagen-related metrics correlated with expert Ashcroft-based severity. In transbronchial biopsy fragments, color-based feature extraction remained feasible, but airway and vascular segmentation was limited by tissue fragmentation and loss of preserved anatomical landmarks. Notably, while this exploratory analysis was not designed to assess clinical outcomes, it demonstrates the feasibility of extending FibroSight from preclinical models to human tissues, while identifying specimen architecture as a key determinant of segmentation performance. Future work should optimize FibroSight for human clinical specimens, validate performance across larger ILD cohorts and biopsy types, and integrate quantitative histological features with radiological, physiological, and clinical outcome data. Such development may support more objective tissue-based phenotyping in translational ILD research and, with appropriate validation, may contribute to future diagnostic pathology workflows.

Taken together, FibroSight provides a scalable and interpretable framework for quantitative histological analysis in ILD. By combining automated structural segmentation with biologically meaningful parameter extraction, the platform enables compartment-resolved characterization of fibrotic, inflammatory, airway, and vascular remodeling within the same tissue section. This multi-parameter approach expands histopathological assessment beyond traditional fibrosis scoring and supports more comprehensive evaluation of disease mechanisms and therapeutic responses in preclinical models. With further validation and methodological refinement, such quantitative digital pathology frameworks may contribute to improved translational research and, ultimately, to more objective histopathological evaluation in human ILD.

## Methods

### Mice

All experiments involving mice conform to the relevant regulatory standards (Technion IACUC and national animal welfare laws, guidelines and policies, approved protocols #IL0830523H for bleomycin and #IL0290224H for influenza).

### Bleomycin induced pulmonary fibrosis

For the induction of pulmonary fibrosis, 10-weeks-old Black 6/c57 mice were gently anesthetized with isoflurane and intubated using a cannula and received 2.25 U/kg of PFI Bleomycin (Baxter) diluted in saline to final volume of 50μL, with saline treated animals serving as controls. Lungs were harvested for 28 days post-bleomycin administration.

### Influenza induced lung tissue damage

To induce pneumonia, Black 6/c57 mice were infected with influenza A virus (A/Puerto Rico/8/1934 (H1N1)) as previously described 39086132. Briefly, viral stocks were grown in embryonated chicken eggs and further propagated in MDCK cells for 3–4 days at 37°C. The viral titer was then determined by plaque assay using MDCK cells. 1.15 × 10³ PFU/mL diluted in PBS ×1 and mice were infected intranasally in a volume of 25 μL. Following infection, mice were monitored daily for clinical signs of pneumonia, including weight loss and respiratory distress. Mice were euthanized between 5–10 days post-infection or upon reaching a humane endpoint of 20% loss of initial body weight, and lungs were harvested for further analysis.

### Lung tissue processing and histology

Tissue samples were fixed in 4% PFA and washed three times with PBS. Following fixation, samples underwent routine dehydration and clearing. Processed tissues were subsequently embedded in paraffin. Paraffin blocks were sectioned at 4μm thickness using a rotary microtome. Tissue sections were deparaffinized and rehydrated and Collagen staining was performed using Pico-Sirius Red solution for 1 hour. SR Slides were digitalized using a Panoramic Flash 250 digital slide scanner (3DHistech, Budapest, Hungary) in bright-field mode using a x20/0.8 objective lens, pixel size 0.242nm with the extended focus mode to select the sharpest image from 3 focal planes spaced 1μm apart. A more detailed protocol is kindly provided in supplementary methods.

### Histopathological Scoring of Fibrosis

To establish a histopathological reference standard for fibrosis severity, Sirius Red–stained lung sections from the bleomycin-induced lung injury model were independently evaluated by two experienced lung pathologists. A total of 30 lung sections, representing a broad range of fibrosis severities, were included in the analysis. Pathologists were blinded to experimental group allocation and quantitative image analysis results. Fibrosis severity was graded using the Ashcroft scoring system, a semi-quantitative scale ranging from normal lung architecture to advanced fibrosis. For each lung section, multiple representative fields of view were examined, and an Ashcroft score was assigned based on established morphological criteria. Scores were averaged across fields to generate a single score for each image. The final reference Ashcroft score for each sample was calculated as the means of the two independent pathologist assessments.

### Histopathological Scoring of Lung Inflammation

Following histological processing, individual lung lobes were separated and embedded in paraffin to enable lobe-specific evaluation. Two trained investigators independently performed histopathological scoring, and their assessments were averaged to generate the final score for each section. Investigators quantified the percentage of inflamed lung tissue in each lobe by evaluating three anatomical compartments based on inflammatory criteria adapted from previous studies(48,49): (1) alveolar/interstitial features including alveolar filling with proteinaceous fluid, epithelial necrosis, inflammatory cell infiltration, septal thickening, and hemorrhage; (2) perivascular features including lymphocytic cuffing, perivascular inflammation, and edema; and (3) peribronchial/peribronchiolar features including epithelial necrosis, peribronchial infiltration, and bronchiolitis. The aggregate involvement across these three compartments was quantified as a percentage using an adapted scoring system(50): 0, no inflammation; 1, only moderate peribronchial Inflammation; 2, <10% inflamed lung tissue; 3, 10 to 25% inflamed lung tissue; 4, 25 to 50% inflamed lung tissue; 5, >50% inflamed lung tissue. These scores were further aggregated to minimal (<10%), moderate (10-50%), or severe (>50%) inflammation for visualization.

### Lung Tissue Segmentation

Whole-lung tissue segmentation was performed using a deep learning–based approach implemented within FibroSight. Histological lung sections stained with Sirius Red were segmented using a pre-trained YOLOv11 convolutional neural network (CNN) optimized for detection and pixel-level segmentation of lung tissue. The model was trained on 725 manually annotated images and validated using a 70/20/10 train–validation–test split.

Model performance metrics included a box precision of 0.997, box recall of 0.993, mAP50of 0.995, and mAP50-95 of 0.987. Pixel-level segmentation achieved a mask precision of 0.997 and a mask recall of 0.993. The definitions and calculation methods for these performance metrics are described in detail in Odeh et al. 2026(51). Training incorporated data augmentation including random rotations, flips, brightness and exposure variation, Gaussian blur, and salt-and-pepper noise. Early stopping was applied with a patience of 100 epochs.

For analysis, users loaded the trained model and batch-processed histological images via the FibroSight GUI. The module automatically generated binary lung masks, segmented lung images, and visual outputs with bounding box overlays. Total lung area (LA) was calculated from the binary masks and exported to an Excel file for downstream normalization of quantitative parameters.

The architectural configuration of the YOLOv11m-seg model is illustrated in Supplementary Fig. S5, and detailed training parameters and performance metrics are summarized in Supplementary Table S5. (Relevant supplementary figures are S5-10, and table S5,6)

### Blood Vessel and Bronchus Segmentation

Segmentation of blood vessels and bronchi was performed using a dedicated deep learning module implemented in FibroSight. This module employs a pre-trained YOLOv11 convolutional neural network optimized for detection and pixel-level segmentation of vascular and bronchial structures in Sirius Red–stained lung sections. The model was trained on 270 manually annotated images and validated using a 70/20/10 train–validation–test split.

Training incorporated the same preprocessing and data augmentation strategies as used for lung tissue segmentation, expanding the dataset to 1,352 images. Model performance achieved mAP@0.5 scores of 0.806 for blood vessels and 0.925 for bronchi, demonstrating reliable detection across a range of lung injury severities.

For analysis, users loaded the trained model and batch-processed images via the FibroSight graphical user interface. The module automatically generated binary masks for blood vessels and bronchi, saved in separate subdirectories, along with visualization outputs displaying class labels and confidence scores. Morphological features were extracted for each segmented structure, including area, centroid, orientation, major and minor axis lengths, eccentricity, equivalent diameter, solidity, and extent. A representative snapshot of this output is provided in Supplementary Table S3. All measurements were exported to an Excel file (“Blood vessels & Bronchus area.xlsx”) for subsequent quantitative analyses.

(Relevant supplementary figures are, S11-15, and table S3, 7-8)

### Analysis of Nuclear Area Fraction

Cellularity analysis was performed using the Nuclei Analysis module within FibroSight. Nuclear segmentation was implemented using intensity thresholding applied to the Value (V) channel of the Hue–Saturation–Value (HSV) color space following RGB conversion. This approach exploits the distinct intensity profile of nuclei in Sirius Red–stained sections while maintaining computational efficiency. Representative images were used to define nucleus-specific lower and upper intensity thresholds in the V channel, which were iteratively optimized using real-time binary mask visualization to ensure accurate segmentation. Once optimized, these thresholds were applied in batch mode to entire image folders.

To account for clustered nuclei and enable estimation of nuclei count, a user-defined division factor corresponding to the independently measured mean nuclear area was specified. Total nuclear area (TNA) was calculated from the binary mask and divided by the division factor to estimate nuclei count (NC). The module automatically generated binary nuclear masks for each image and exported quantitative outputs, including TNA and NC, to an Excel file (“Number of nuclei.xlsx”). Cellularity metrics were subsequently derived by normalizing TNA to total lung area.

### Collagen and Tissue extraction

Collagen and tissue compartments were segmented using LAB color space conversion with thresholding on the A channel (green–red axis). Lower and upper intensity thresholds were defined on representative images and subsequently applied in batch mode across all samples within each experimental set.

Binary masks were generated for collagen and total tissue. Collagen area (CA) and tissue area (TA) were calculated as the total segmented pixel counts per image. Pixel-wise overlap analysis between collagen and parenchyma masks enabled compartment-specific quantification. Parenchymal collagen (PC) was defined as collagen excluding vascular and bronchial overlap regions. Parenchymal tissue area (PTA) excluded vascular and bronchial wall regions. Clean lung area (CLA) was defined as lung area minus vascular and bronchial areas.

### Definition and Derivation of Quantitative Parameters

Quantitative parameters were derived from primary area measurements obtained from the lung segmentation, blood vessel and bronchus segmentation, nuclei analysis, and collagen/parenchyma modules. Primary measurements included lung area (LA), total tissue area (TA), collagen area (CA), bronchi area (BR), blood vessel area (BV), total nuclear area (TNA), and region-specific overlap areas including perivascular collagen (BVC), peribronchial collagen (BRC), vascular wall tissue (BVT), and bronchial wall tissue (BRT). To isolate the functional gas-exchange compartment, derived anatomical regions were defined as follows: clean lung area (CLA) was calculated as lung area excluding bronchi and blood vessels (CLA = LA − [BV + BR]). Parenchymal tissue area (PTA) was defined as total tissue area excluding vascular and bronchial wall components (PTA = TA − [BVT + BRT]). Parenchymal collagen (PC) was calculated as total collagen area minus collagen overlapping with blood vessels and bronchi (PC = CA − [BVC + BRC]). Parenchymal Fibrosis parameter was generated by normalizing parenchymal collagen per clean lung area (PC/CLA). Parenchymal tissue density was calculated as (PTA/ CLA) and the complementary value of Parenchymal airspace fraction was calculated as (CLA − PTA)/CLA. Cellularity metrics were derived from total nuclear area and estimated nuclei count, normalized to lung area (TNA/LA). Region-specific remodeling parameters were calculated to quantify compartmental fibrosis and wall thickening, including: vascular wall area fraction (BVT/BV), bronchial wall area fraction (BRT/BR), and the optional perivascular fibrosis (BVC/BV), peribronchial fibrosis (BRC/BR) parameters.

A complete list of primary outputs, definitions, and abbreviations is provided in Supplementary Table S2.

### Image Aggregation and Per-Animal Quantification

For validation against expert histological scoring, quantitative parameters were calculated per image, with each Sirius Red–stained lung lobe analyzed independently and compared to its corresponding integrated Ashcroft score. For animal model experiments, quantitative outputs from multiple lung lobes of the same mouse were aggregated to generate a single per-animal value. Aggregation was performed using area-weighted averaging based on the segmented lung area of each lobe, followed by normalization to the relevant summed areas (e.g., lung area, tissue area, or clean lung area). Statistical comparisons were performed using these per-animal values.

### Human ILD biopsy analysis

Human lung biopsy specimens were obtained retrospectively from patients evaluated for interstitial lung disease at Rambam Health Care Campus, Haifa, Israel. The study was approved by the institutional review board/Helsinki Committee (approval number RMB-D-0504-24), and all procedures were conducted in accordance with the Declaration of Helsinki. Patient data and tissue samples were anonymized prior to analysis. Formalin-fixed, paraffin-embedded sections were stained with Sirius Red for FibroSight analysis, with corresponding H&E-stained sections used for histopathological reference. Available clinical and diagnostic data were reviewed for specimen classification.

Two specimen types were included: surgical wedge-resection specimens and transbronchial biopsies (TBB). Wedge-resection specimens were obtained from three patients and represented 12 paraffin blocks in total: one block from patient 1, seven blocks from patient 2, and four blocks from patient 3. Because wedge specimens contain large and heterogeneous tissue areas, digitized slides were divided into non-overlapping, anatomically coherent tissue regions comparable in scale to murine lung lobe images, yielding 32 image-level regions. Each region was assigned an independent Ashcroft-based fibrosis score by manual histopathological assessment, blinded to FibroSight outputs. FibroSight was applied to the corresponding Sirius Red regions to extract tissue, collagen, nuclear, airway, and vascular features. Automated tissue, blood vessel, and bronchial masks were reviewed and manually curated when necessary, including correction of missed microvascular structures and removal of falsely segmented bronchial regions.

TBB specimens were obtained from 16 patients, represented by 16 paraffin blocks. Digitized slides were cropped into tissue-containing regions suitable for analysis, yielding 48 cropped tissue-region images. Because TBB specimens are small and frequently affected by tissue compression, fragmentation, and reduced preservation of anatomical landmarks, TBB analysis was restricted to tissue segmentation and color-based extraction of collagen, tissue, and nuclear features. Automated tissue segmentation performed reliably on cropped TBB regions; however, blood vessel and bronchial segmentation was inconsistent and was therefore excluded from downstream TBB analyses.

All human biopsy analyses were performed at the image-region level. Because multiple wedge-derived regions originated from the same patients and paraffin blocks, these analyses were considered exploratory and intended to assess technical and translational feasibility rather than patient-level clinical validation.

### Statistical Analysis

Statistical analyses were performed using MATLAB (MathWorks). Data is presented as mean ± standard error of the mean (SEM), unless otherwise stated. Each data point represents a single animal (for murine models) or a single lung lobe image (for Ashcroft validation analysis), as specified in the corresponding Results section. Normality of data distribution was assessed using the Shapiro–Wilk test. For comparisons between two independent groups, normally distributed data were analyzed using unpaired two-tailed Student’s t-test, while non-normally distributed data were analyzed using the Mann–Whitney U test. Correlations between FibroSight-derived parameters and Ashcroft scores were evaluated using Spearman’s rank correlation coefficient (ρ), given the ordinal nature of Ashcroft scoring. For analysis of blood vessel size and morphology distributions, the Kolmogorov–Smirnov test was used to compare cumulative distributions between experimental groups. P-value < 0.05 was considered statistically significant. Gemini 2.5 Flash (Google) was used to assist in drafting and debugging MATLAB scripts for data processing, plotting, and statistical analyses. All scripts and outputs were reviewed and verified by the authors.

### Software Development and AI Disclosure

The core source code for the software was developed entirely by the authors without the assistance of AI tools. The generative AI model Gemini 2.5 Flash (Google) was utilized to assist in refining the design and layout of the user interface. All AI-assisted interface components were manually reviewed and verified prior to implementation.

## Data and code availability

The source code, software tools, user manual, and underlying data supporting the findings of this study are openly available in the FibroSight GitHub repository at: https://github.com/Anas-Odeh/FibroSight. To ensure broad accessibility for researchers without extensive computational backgrounds, FibroSight is provided as a standalone Windows executable (.exe), enabling one-click installation and use with minimal configuration. A detailed user manual is provided to guide installation, workflow execution, threshold calibration, batch analysis, and interpretation of software outputs. Further updates, version releases, and extended documentation will be maintained within the same repository.

## Acknowledgements

We would like to thank Maya Holdengreber and Melia Gurewitz from the Technion Biomedical Core Facility unit for assistance with the Automatic slide scanner. We would like to thank Katren Sakran for the histology service. Yuanning Guo gratefully acknowledges Yuanxi Zeng, Taoyu Wang, Tianjiao Li, and Zhicheng Huang for their insightful discussions and collaborative exploration of histological scoring in prior studies of mouse radiation-induced pneumonitis and fibrosis models, which provided important experience and guidance for Yuanning Guo’s tissue assessment and segmentation work in this study. ChatGPT, GPT-5.5 Thinking (OpenAI), was used as a writing assistant to help revise author-written manuscript drafts for grammar, readability, clarity, and consistency. The initial drafts of all manuscript sections were written by the authors, and all AI-assisted suggestions were reviewed, edited, and approved by the authors. The authors take full responsibility for the final manuscript content.

## Author Contributions

I.M. and A.O. conceived the study. I.M. led the biological domain of the project, performed the in vivo bleomycin mouse experiments, conducted the biological analyses, and wrote the original manuscript draft. A.O. led the computational domain of the project, conceptualizing the technical framework and developing the FibroSight software platform, generating the computational figures and tables, and contributing to manuscript writing and reviewing. Y.G. contributed to image annotation, data organization, figure preparation, and manuscript writing. J.H. contributed to image annotation. N.M. contributed to image annotation. M.A. and T.S. assisted with mouse experiments and tissue collection. A.R. performed the influenza infection experiments and tissue collection. M.H. and H.A.C. performed inflammatory scoring of influenza-infected mice. R.P designed and supervised the influenza model component and contributed to interpretation of the inflammatory remodeling data. R.B.S. collected and organized the human biopsy data and contributed to image annotation. Y.D. contributed to the human biopsy component and manuscript preparation. I.L. and P.S. performed expert Ashcroft scoring. A.O, A.S, H.W. and P.H. conceptualized the overarching research goals, supervised the project, and contributed to manuscript review and editing.

## Supplementary Material

**Table S1A.** Overview of existing computational tools for histological fibrosis quantification.

| Study | Year | Stain | Automation level | Accessibility |
| --- | --- | --- | --- | --- |
| Gilhodes et al. | 2017 | Masson's Trichrome | Automated quantification with manual exclusion of large bronchi and vessels | Proprietary software; not publicly accessible |
| Seger et al. | 2018 | Masson's Trichrome | Fully automated SVM-based segmentation using Orbit | Requires Orbit; plugin available but limited documentation |
| Heinemann et al. | 2018 | Masson's Trichrome | CNN-based analysis after manual selection of alveolar tiles | No public GUI; method described in publication |
| Ségard et al. | 2024 | Sirius Red and Masson's Trichrome | Semi-automated ImageJ macro toolbox | Open-source ImageJ/Fiji plugin with documentation |
| Goto et al. | 2025 | Sirius Red | Fully automated CNN workflow using pre-selected fibrosis-rich patches | Published methodology only; no public tool release |
| Drakeley et al. | 2025 | H&E; Trichrome | Semi-automated tissue segmentation with automated thresholding | Python-based GUI; requires CUDA-enabled GPU and technical setup |

Abbreviations: CNN, convolutional neural network; CUDA, Compute Unified Device Architecture; GUI, graphical user interface; SVM, support vector machine.

**Table S1B.**
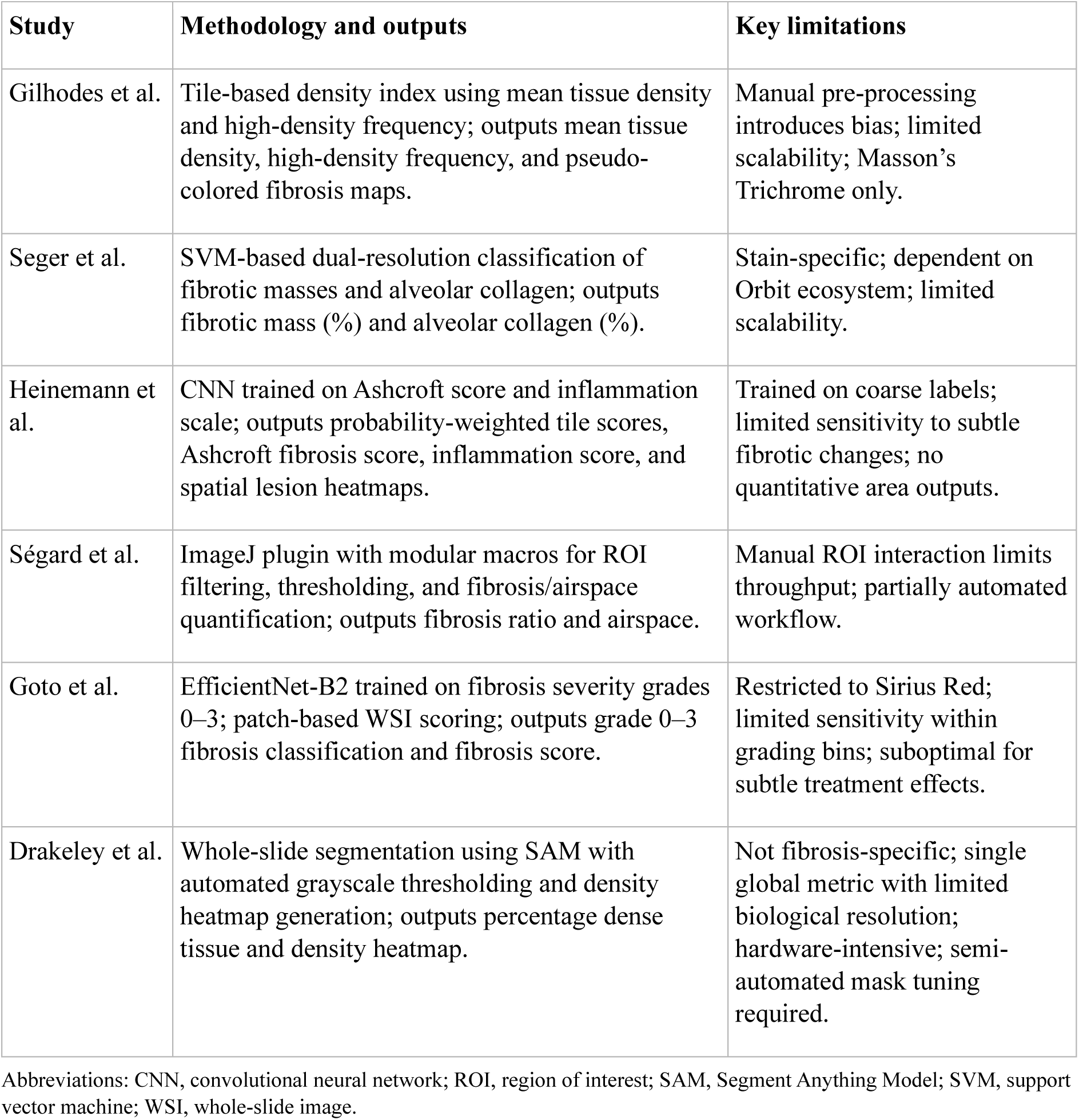
Methodology, outputs, and limitations of existing tools.

**Table S2.**
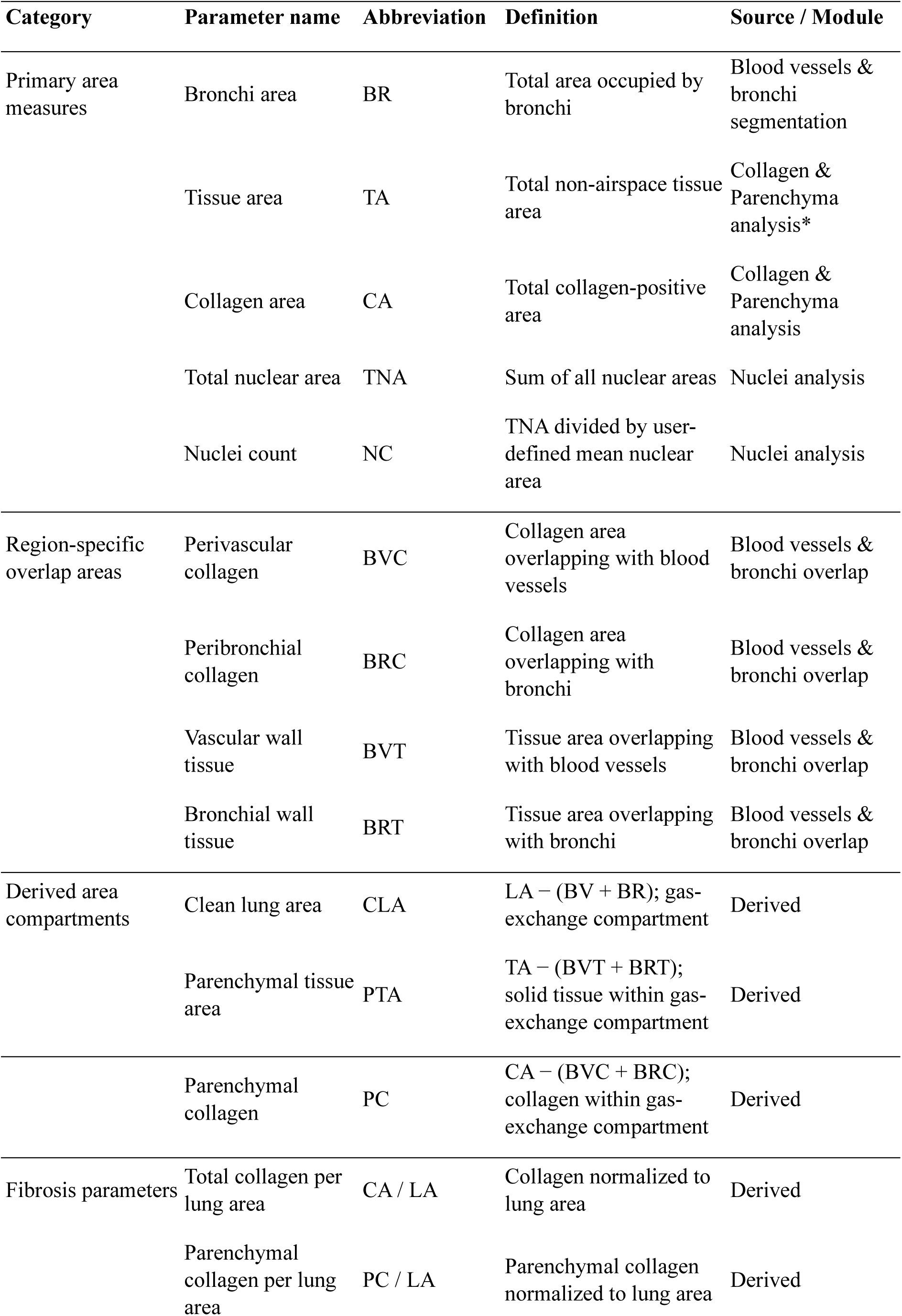
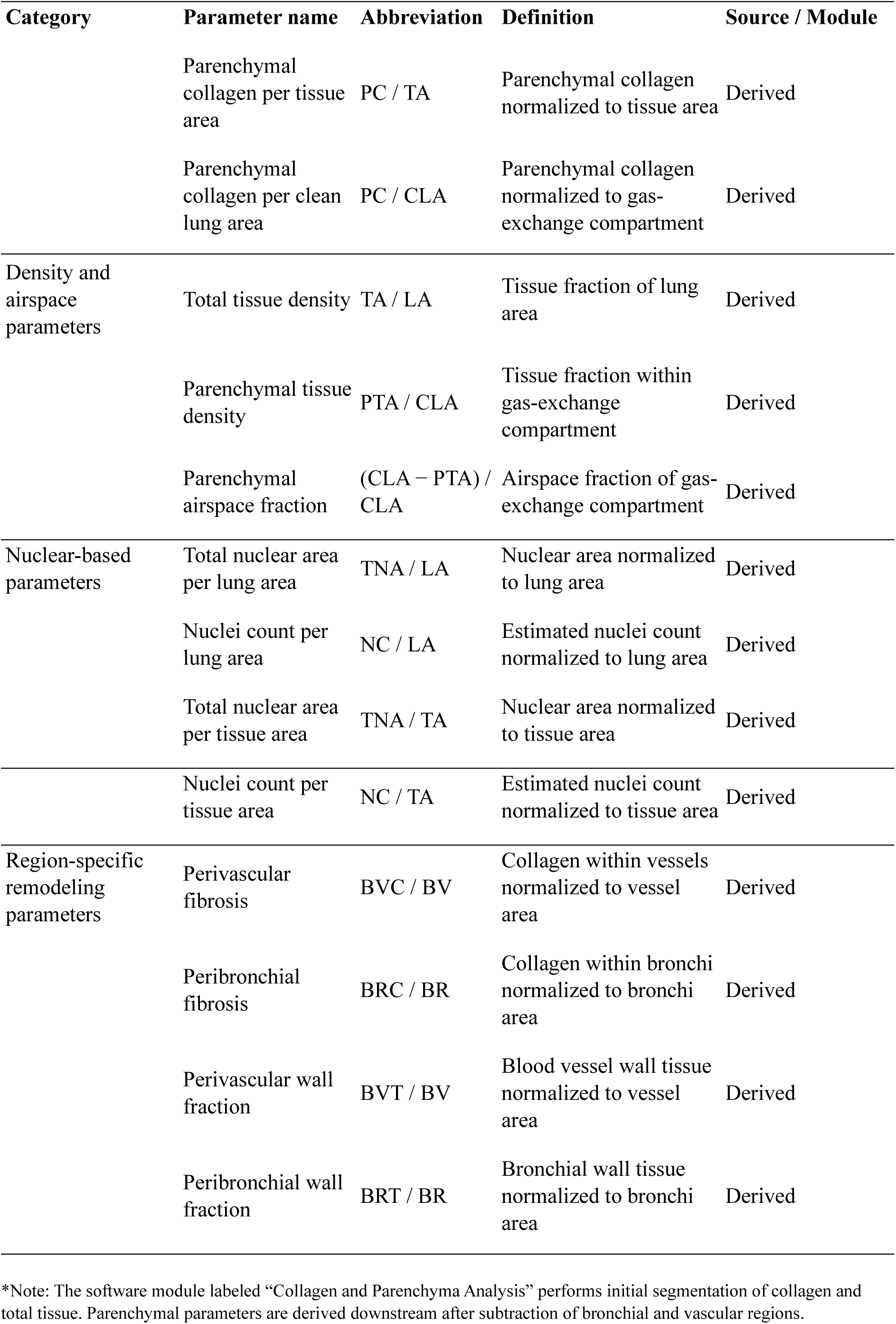
FibroSight area outputs and derived quantitative parameters.

**Figure S1.**
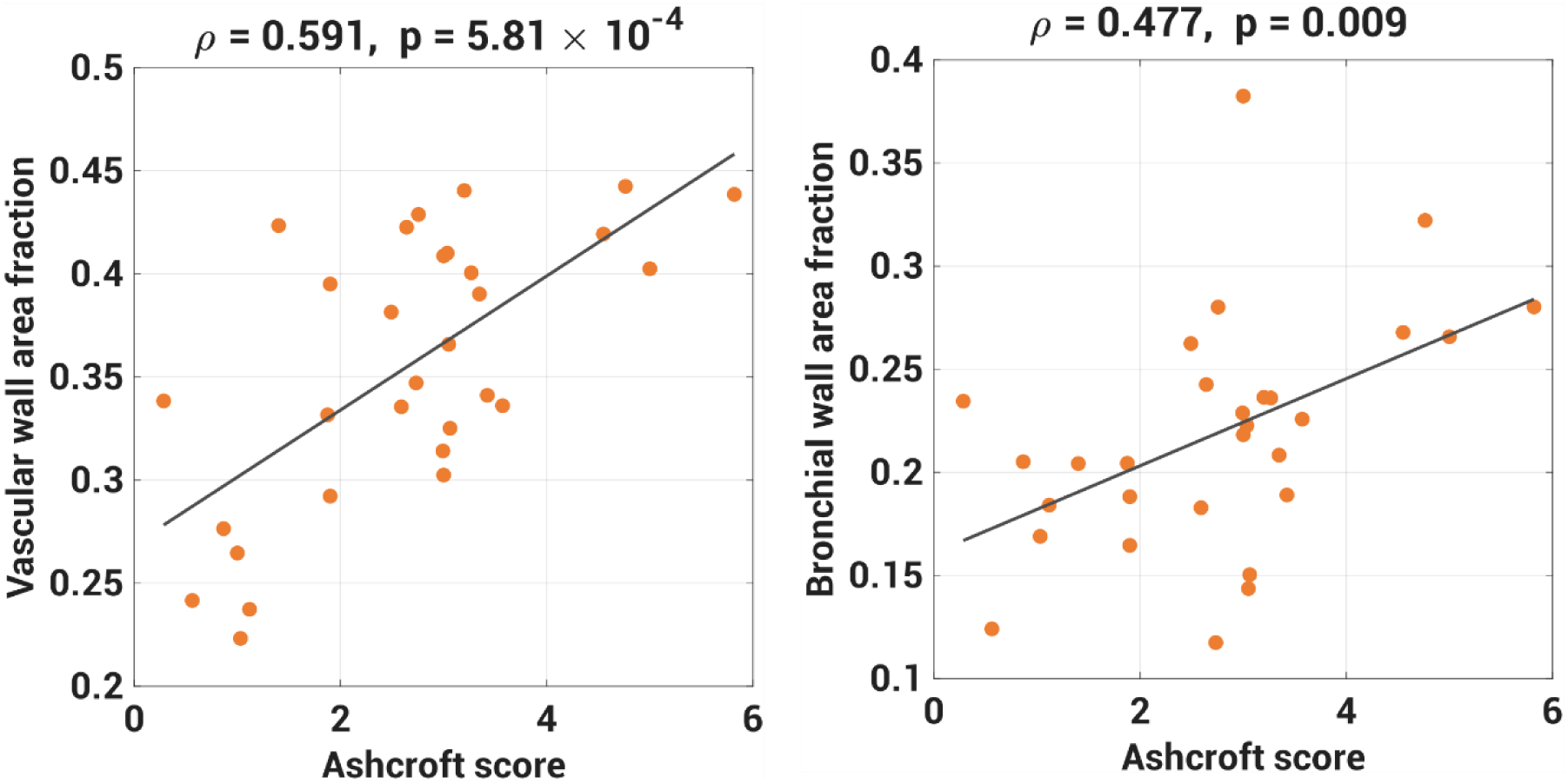
Association of airway and vascular wall remodeling with histopathological fibrosis severity. **(A)** Correlation between vascular wall area fraction and Ashcroft score (ρ = 0.591, p = 5.81 × 10⁻⁴). **(B)** Correlation between bronchial wall area fraction and Ashcroft score (ρ = 0.477, p = 0.0089). Each dot represents a Sirius Red-stained lung section from the bleomycin model. Wall area fraction was calculated as wall tissue area normalized to total vessel or bronchial area, respectively. Both parameters demonstrate moderate positive correlations with increasing fibrosis severity. Spearman correlation coefficients (ρ) and corresponding p values are shown.

**Table S3.**
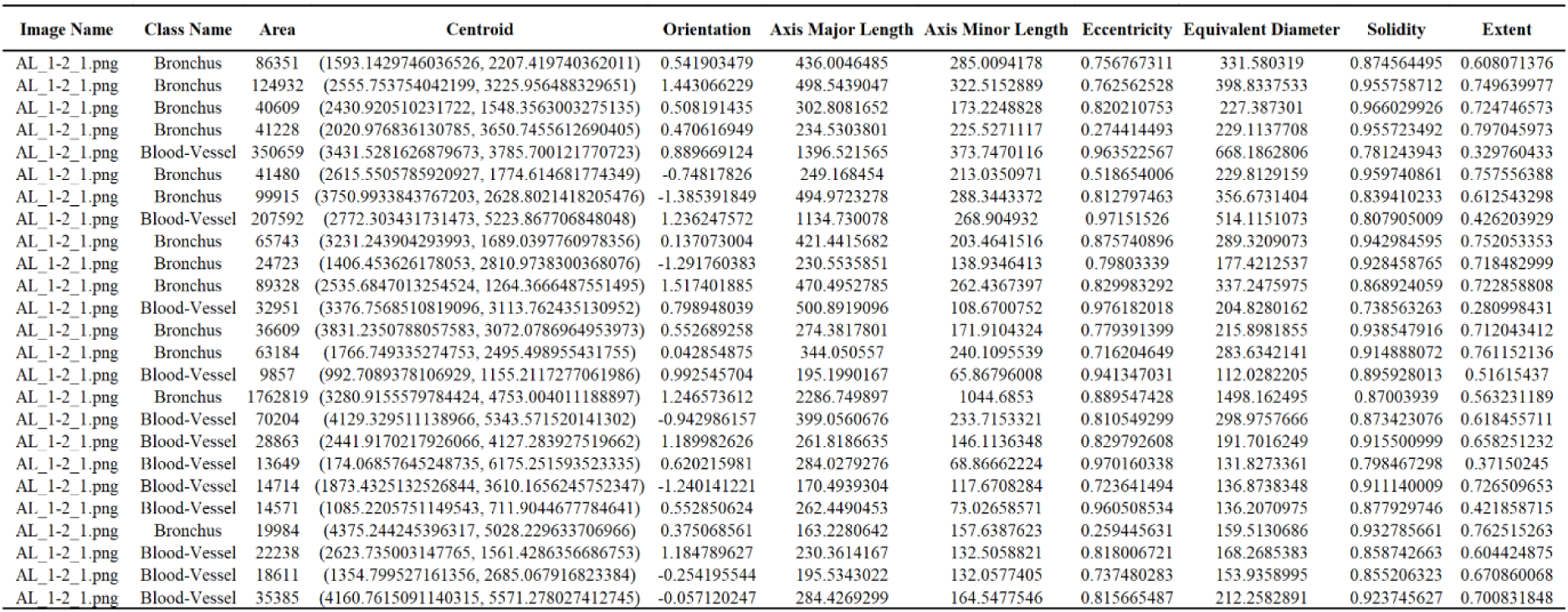
A Snapshot of Detailed Morphometric Analysis of Segmented Vascular and Bronchial Structures. Quantitative morphological data derived from the automated segmentation of blood vessels and bronchi in lung histological sections. The parameters include geometric properties (Area, Axis Lengths), spatial location (Centroid Coordinates), and shape descriptors (Orientation, Eccentricity, Extent, Solidity). These metrics provide a detailed, object-specific analysis of each segmented structure, enabling a precise characterization of lung tissue microarchitecture. Note that the Image Name column may contain multiple entries. This indicates that each histological image can contain multiple distinct blood vessels or bronchi, which are independently segmented and quantified. Consequently, each row corresponding to a repeated Image Name represents a unique vascular or bronchial element within that image, facilitating a detailed, object-level assessment of lung tissue morphology.

**Figure S2.**
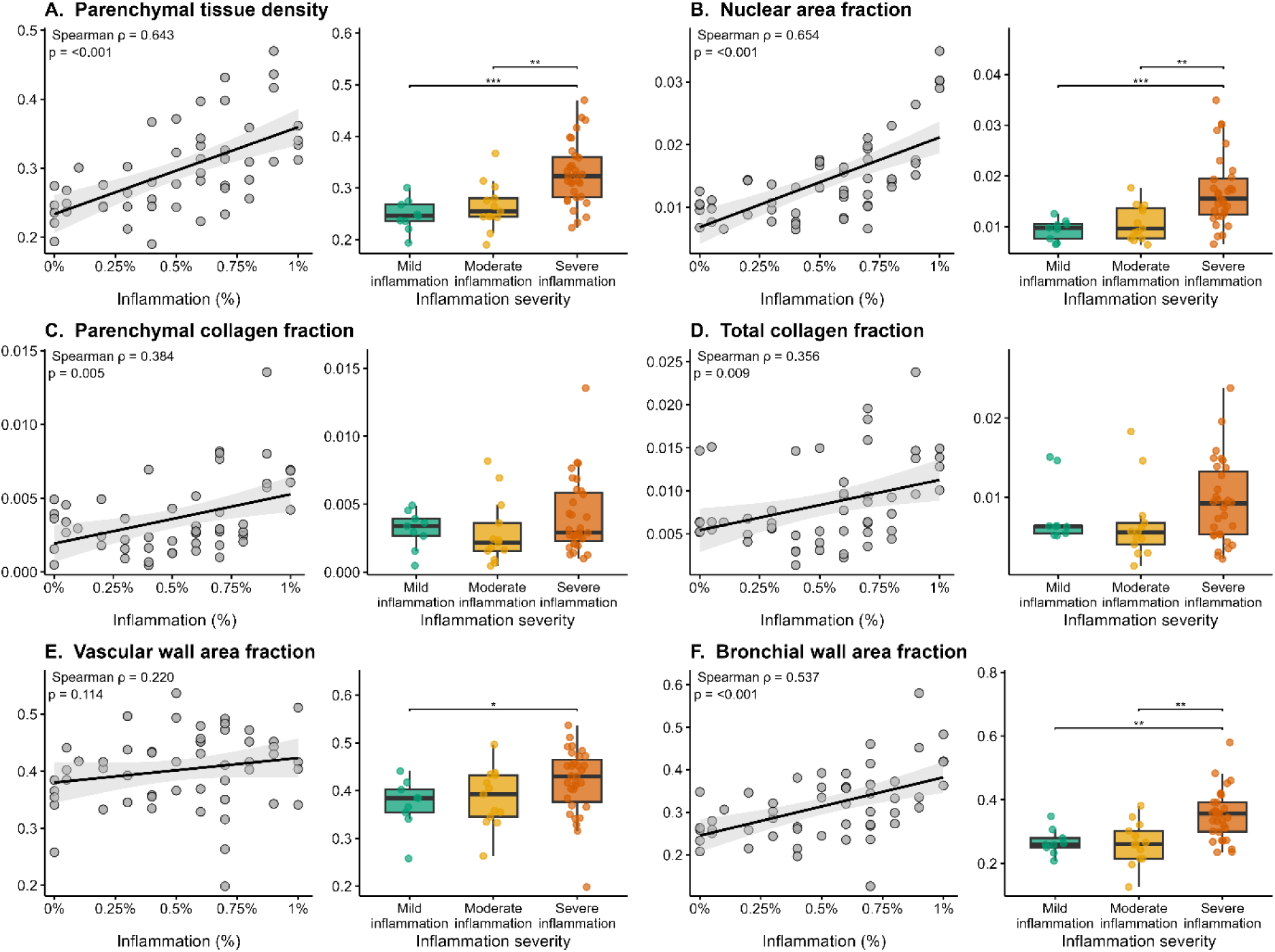
Correlation between FibroSight-derived metrics and semi-quantitative inflammatory grading. FibroSight-derived lobe-level parameters were compared with an adapted semi-quantitative inflammatory grading system in the influenza-induced lung injury model. For each parameter, the left plot shows correlation with inflammatory involvement scored as percentage of affected lung tissue, and the right plot shows severity-stratified analysis across mild, moderate, and severe inflammation groups. **(A)** Parenchymal tissue density correlated positively with inflammatory grading (ρ = 0.643, p < 0.001) and was increased in severely inflamed lobes. **(B)** Nuclear area fraction showed a strong positive correlation with inflammatory grading (ρ = 0.654, p < 0.001), consistent with increased cellular infiltration, and was elevated in severe inflammation. **(C, D)** Parenchymal collagen fraction and total collagen fraction showed weaker but significant positive correlations with inflammatory grading (ρ = 0.384, p = 0.005 and ρ = 0.356, p = 0.009, respectively). **(E)** Vascular wall area fraction was not significantly correlated with inflammatory grading (ρ = 0.220, p = 0.114) but was increased in severely inflamed lobes. **(F)** Bronchial wall area fraction correlated positively with inflammatory grading (ρ = 0.537, p < 0.001) and was increased in severely inflamed lobes. Correlations were calculated using Spearman’s rank correlation. Severity-group comparisons were performed using pairwise Wilcoxon tests with Benjamini– Hochberg correction. Each point represents an individual lung lobe.

**Table S4.** Manual curation log for human wedge-resection biopsy mask correction. Human wedge-resection tissue-region images were reviewed following automated FibroSight segmentation. When necessary, blood vessel and bronchial masks were manually curated to correct missed structures or false-positive segmentations before downstream compartment-specific quantification. Each row represents one analyzed tissue-region image. BV, blood vessel; BR, bronchus.

| **Tissue region** | **Visual evaluation after automated segmentation** | **Manual correction** |
| --- | --- | --- |
| 1 | No correction required | No correction |
| 2 | Large bronchi missing | Added 2 large bronchi and 1 large vessel |
| 3 | Large bronchi missing | Added 2 large bronchi |
| 4 | One BV and one bronchus missing | Added 1 large vessel |
| 5 | No correction required | No correction |
| 6 | No correction required | No correction |
| 7 | No correction required | No correction |
| 8 | One BV missing | Added 1 BV |
| 9 | No correction required | No correction |
| 10 | No correction required | No correction |
| 11 | No correction required | No correction |
| 12 | No correction required | No correction |
| 13 | One large BV missing | Added 2 BVs |
| 14 | Medium-sized BV missing | Added 1 BV |
| 15 | Large BV and bronchi missing | Added 2 BVs and 1 BR |
| 16 | One bronchus missing | No correction documented |
| 17 | Some medium-sized bronchi missing | No correction documented |
| 18 | Some small BVs missing | No correction documented |
| 19 | Incomplete blood vessel wall segmentation and one medium-sized bronchus missing | No correction documented |
| 20 | Large blood vessels missing after correction | Added 2 BVs and deleted 1 false-positive mask |
| 21 | Some bronchi missing | Added 3 bronchi and 2 vessels |
| 22 | No correction required | No correction |
| 23 | Medium/small BVs missing | Added 2 BVs and 1 BR |
| 24 | No correction required | No correction |
| 25 | No correction required | No correction |
| 26 | Medium-sized bronchi missing | Added 1 BV and 2 BRs |
| 27 | No correction required | No correction |
| 28 | Major BV missing | Added 2 BVs |
| 29 | No missing structure documented | Added 1 BV and 1 BR |
| 30 | Medium-sized BV and bronchus missing | No correction documented |
| 31 | No correction required | No correction |
| 32 | No correction required | No correction |

**Figure S3.**
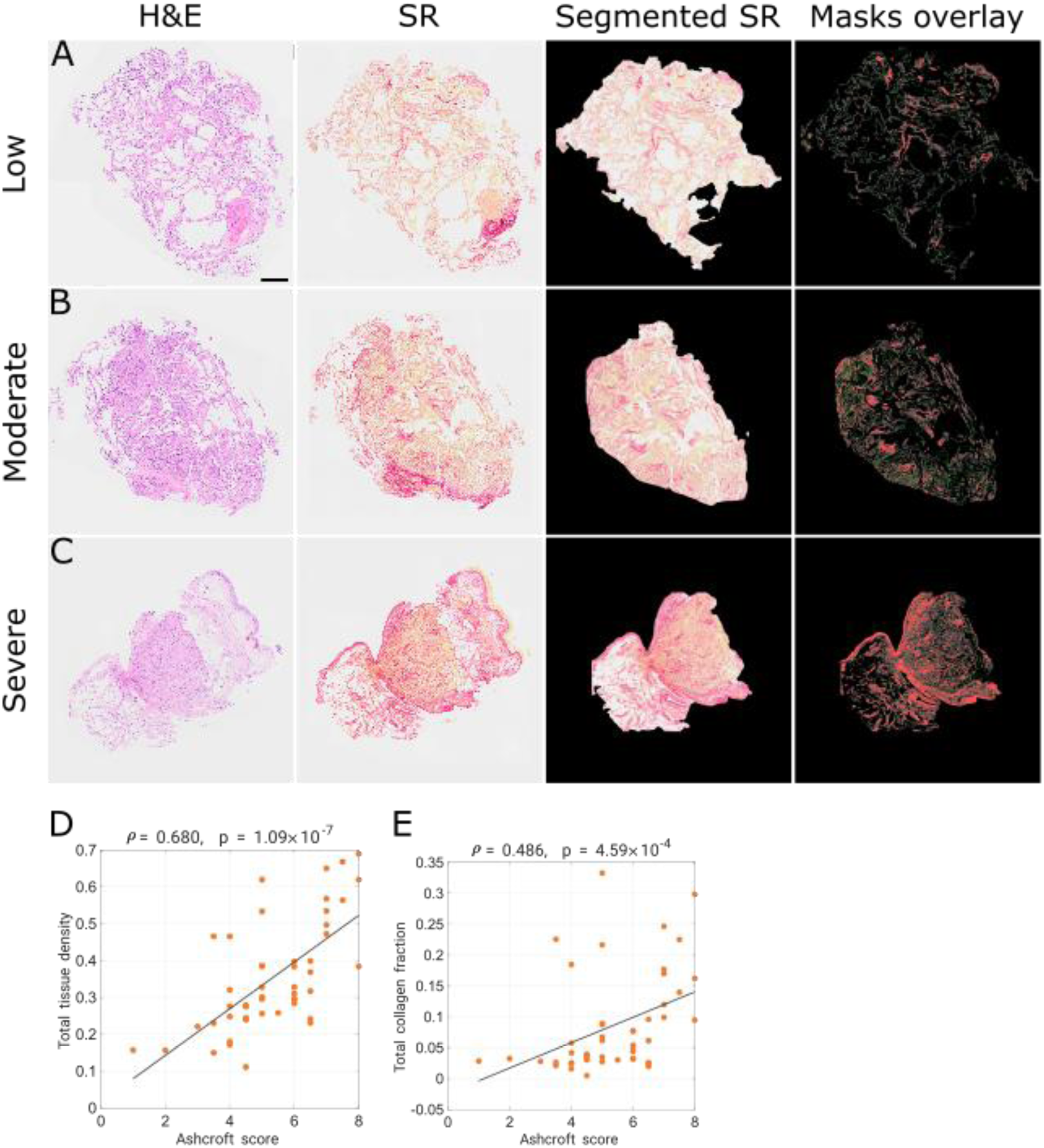
FibroSight enables tissue-level quantitative feature extraction from human transbronchial biopsy specimens. **(A–C)** Representative human transbronchial biopsy tissue regions with low, moderate, and severe histological remodeling. For each region, H&E-stained images, matched Sirius Red (SR)-stained images, segmented SR regions, and FibroSight-derived mask overlays are shown. SR input regions were manually curated before analysis to exclude non-parenchymal structures, including large bronchial tissue. FibroSight subsequently delineated tissue borders within the curated biopsy regions and generated masks for tissue, collagen, and nuclei. In the mask overlays, gray indicates tissue, red indicates collagen, and green indicates nuclei. Blood vessel and bronchus masks are not shown because automated vascular and airway segmentation was not reliable in transbronchial biopsy specimens. **(D, E)** Correlations between expert Ashcroft-based fibrosis scores and FibroSight-derived tissue-level parameters in transbronchial biopsy regions. Total tissue density correlated with Ashcroft score (ρ = 0.680, p = 1.09 × 10⁻⁷; D), and total collagen fraction showed a more moderate positive correlation with Ashcroft score (ρ = 0.486, p = 4.59 × 10⁻⁴; E). Each point represents one analyzed biopsy region. Correlations were calculated using Spearman’s rank correlation.

**Figure S4.**
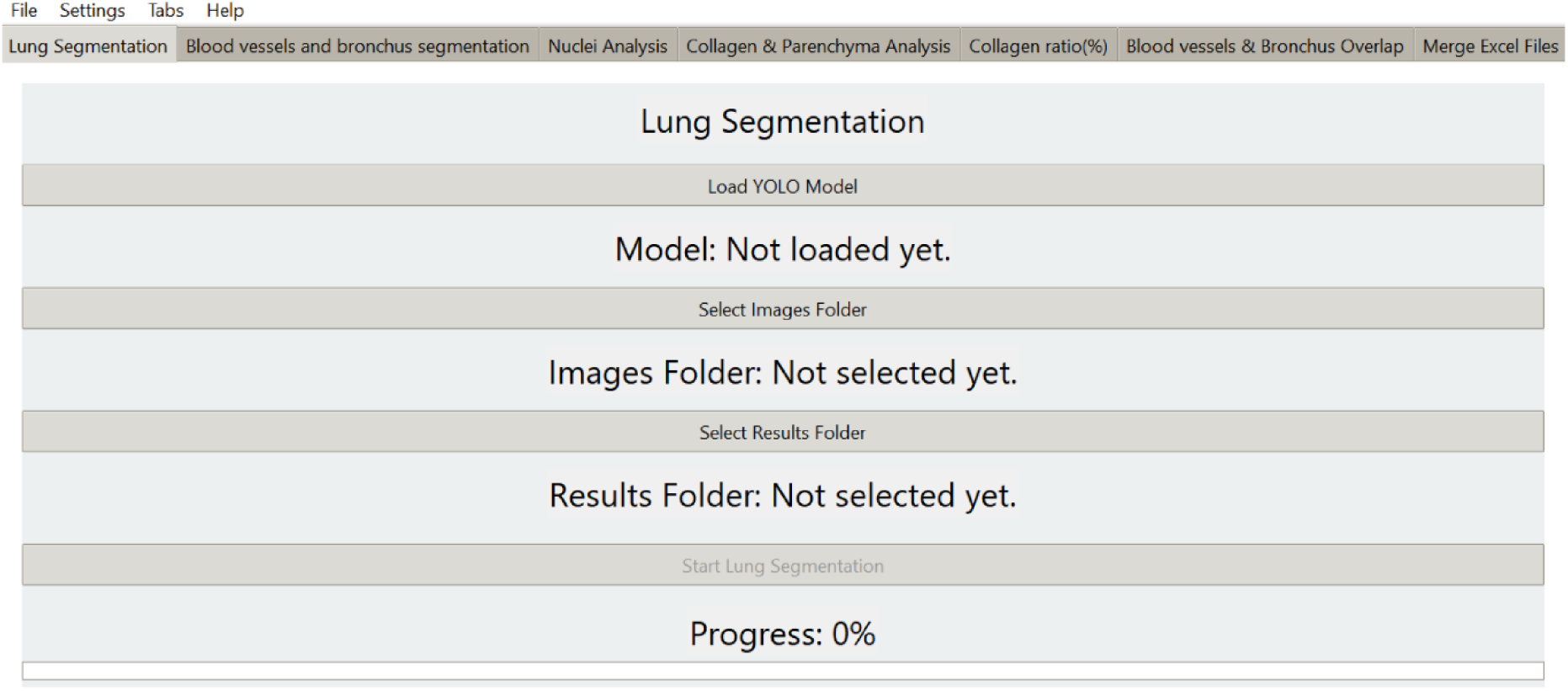
FibroSight’s Modular Workflow for Lung Fibrosis Analysis. FibroSight interface is organized into a series of modular tabs, each dedicated to a specific step in the lung fibrosis analysis workflow. The Lung Segmentation tab allows users to load a YOLO model, select image and results folders, and initiate automated lung segmentation. Subsequent tabs provide tools for blood vessel and bronchus segmentation, nuclei analysis, collagen and parenchyma analysis, collagen ratio quantification, blood vessel and bronchus overlap analysis, and merging of Excel data files.

**Figure S5.**
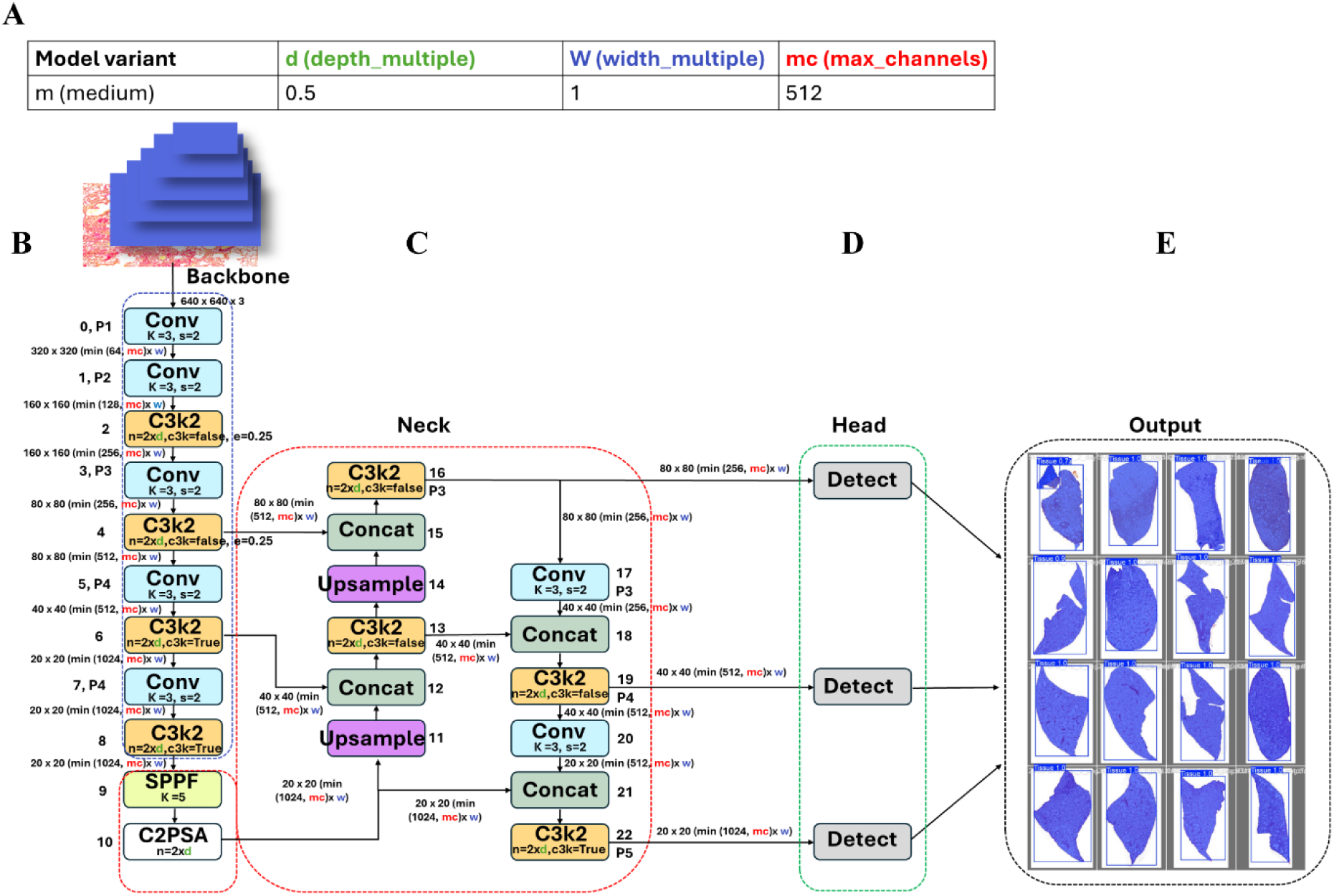
Architecture of the YOLOv11m-seg Model within FibroSight for Lung Tissue Segmentation. **(A)** A table outlines the configuration parameters for the medium (m) variant of the YOLOv11m-seg model used in FibroSight, including depth multiplier (d), width multiplier (w), and maximum channel count (mc). **(B)** Inputting the lung tissue image into a backbone network, which utilizes convolutional layers and C3K2 modules to extract multi-scale features. **(C)** The neck section of the network combines these multi-scale features through concatenation and up sampling operations, creating a unified feature representation. **(D)** The head component performs object detection and segmentation at three distinct scales, generating the final lung tissue segmentation masks. **(E)** An example output grid demonstrates the model’s ability to accurately segment lung tissue regions as processed by FibroSight.

**Figure S6.**
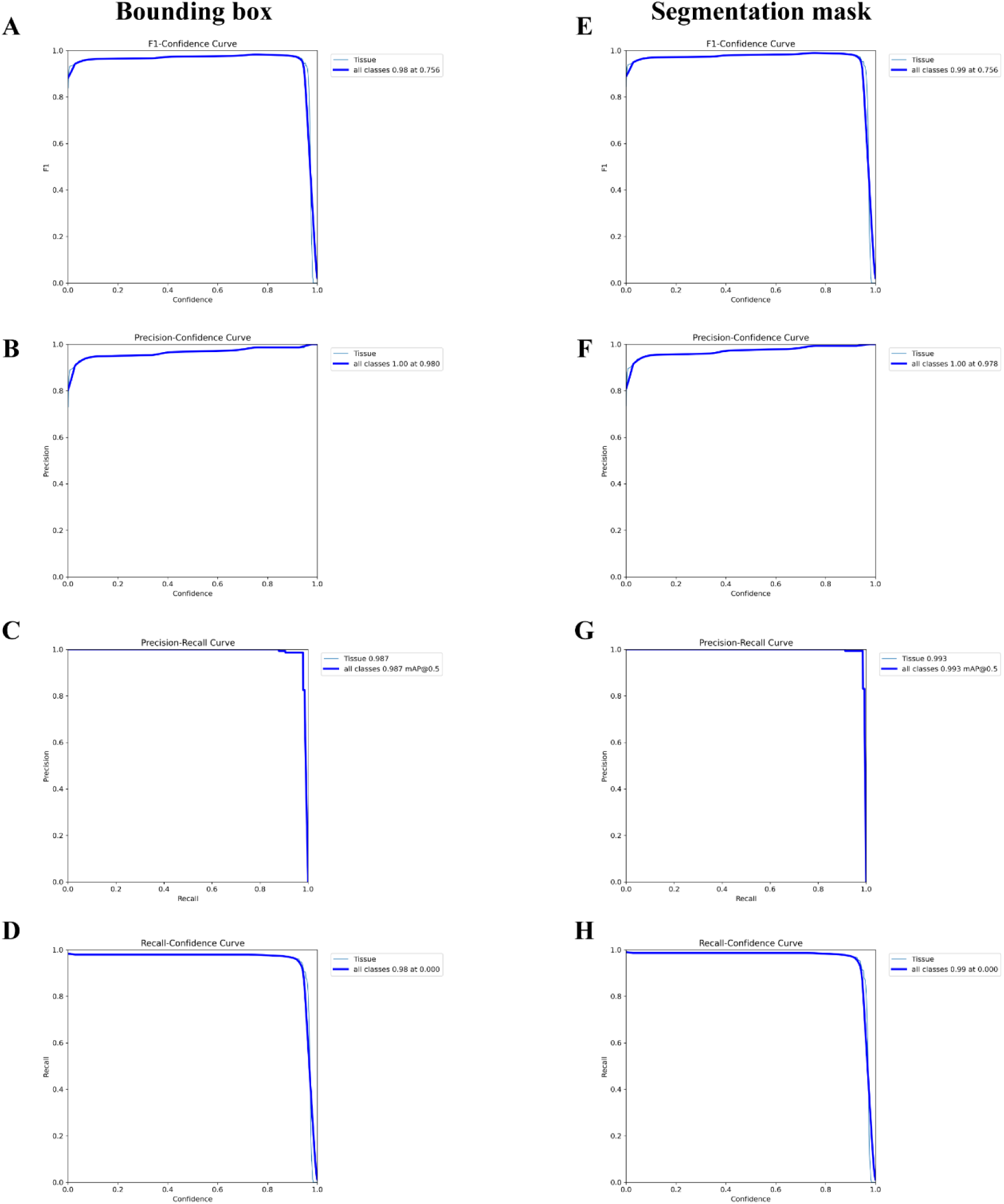
Performance Evaluation Curves for YOLOv11m-seg Lung Tissue Segmentation. **(A-H)** A comprehensive performance evaluation curves for the YOLOv11m-seg model’s lung tissue segmentation, illustrating the relationship between confidence thresholds and key performance metrics. **(A-D)** Illustrate the bounding box prediction performance, while panels **(E-H)** illustrate segmentation mask performance. Specifically, **(A)** and **(E)** show the F1-Confidence curves, with the maximum F1 score of 0.98 and 0.99, respectively. **(B)** and **(F)** show the Precision-Confidence curves, with the precision at maximum confidence of 1.0 for both. **(C)** and **(G)** show the Precision-Recall curves, with precision of 0.987 and 0.993, respectively. **(D)** and **(H)** show the Recall-Confidence curves, with the recall at maximum confidence of 0.98 and 0.99, respectively. In each panel, the blue line represents the Tissue class (lung tissue). These curves provide a comprehensive assessment of the model’s performance sensitivity to confidence thresholds, demonstrating its efficacy in lung tissue detection and segmentation.

**Figure S7.**
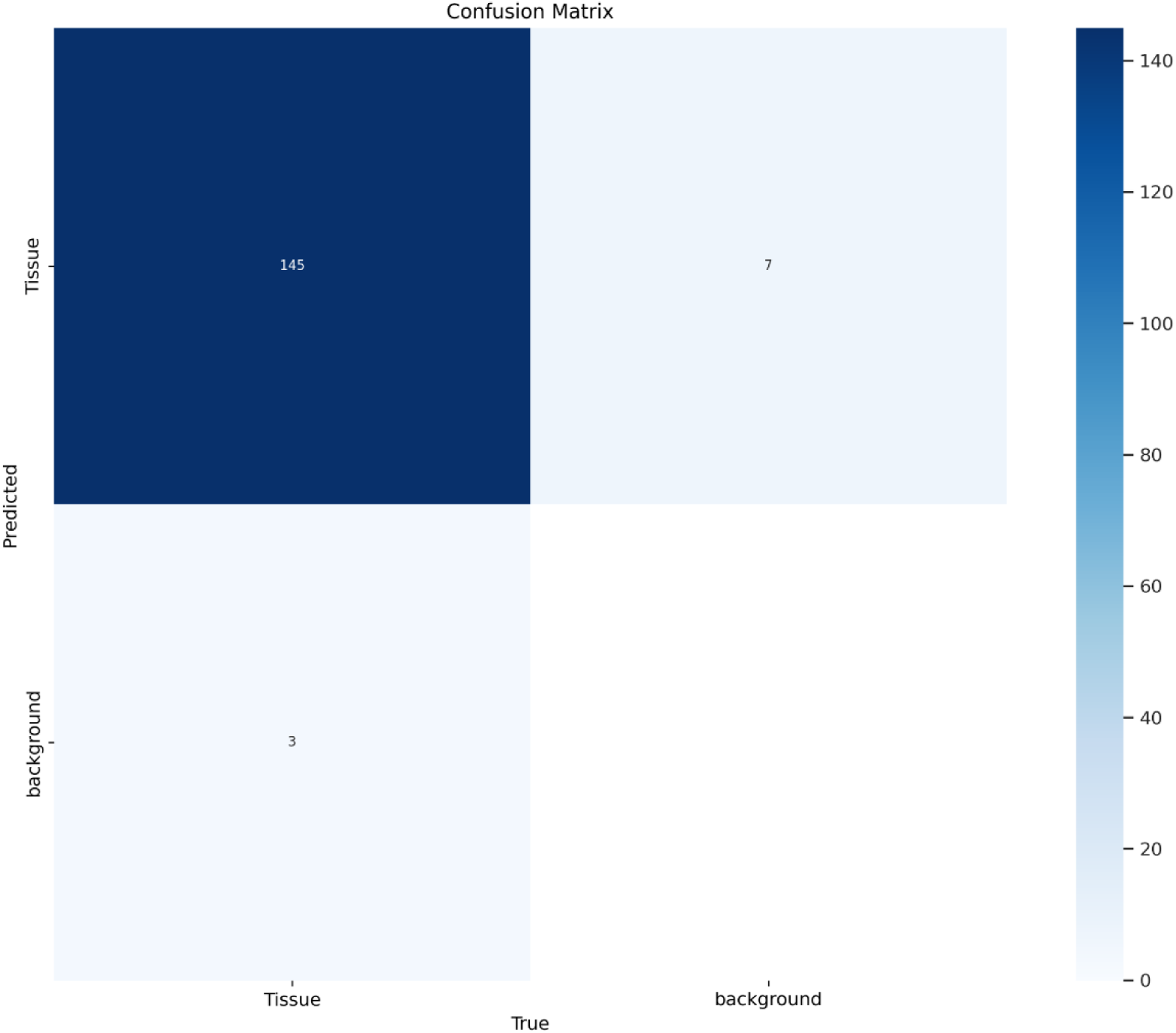
Confusion Matrix for Lung Tissue Segmentation. Confusion matrix for the YOLOv11m-seg model’s lung tissue segmentation performance, displaying the model’s classification accuracy between Tissue (lung tissue) and background. The rows represent the predicted labels, and the columns represent the true (actual) labels. The numerical values within each cell indicate the count of pixels falling into each category. The matrix reveals that 145 pixels were correctly classified as Tissue, and 3 pixels were misclassified as background when they were Tissue. Similarly, 2 pixels were misclassified as Tissue when they were background. The color bar on the right represents the count scale, with darker shades indicating higher pixel counts.

**Figure S8.**
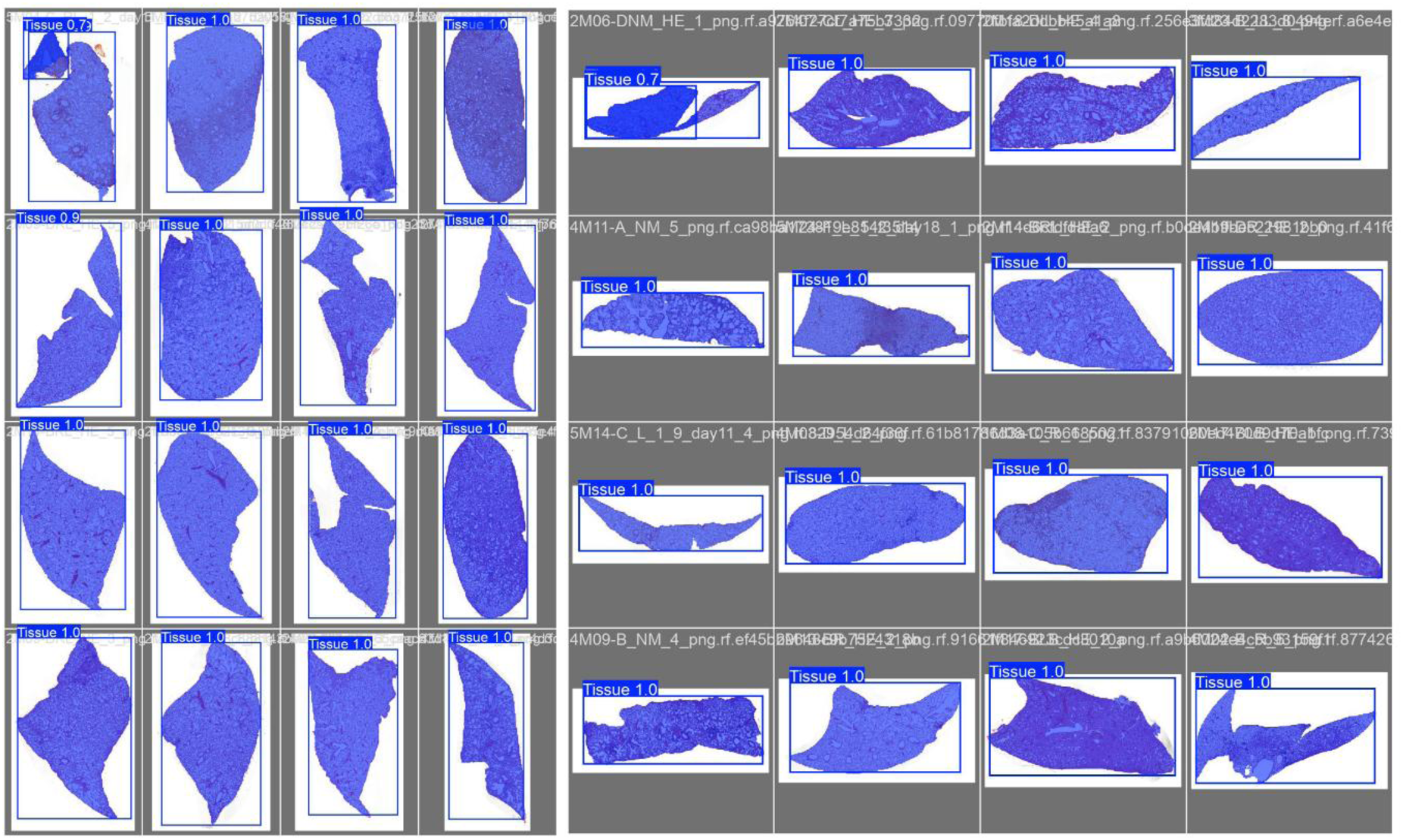
YOLOv11m-seg Validation Results of Lung tissue Segmentation. Batch of sample lung tissue segmentation results generated by the YOLOv11m-seg model. Each image in the grid shows the original histological lung tissue image with an overlaid blue bounding box and segmentation mask, highlighting the detected lung tissue region. The Tissue label, along with a confidence score (ranging from 0.79 to 1.0), is displayed above each bounding box. The confidence score reflects the model’s certainty in the segmentation accuracy. These numbers report the model’s calculated certainty and show how it outlines the tissue boundaries.

**Figure S9.**
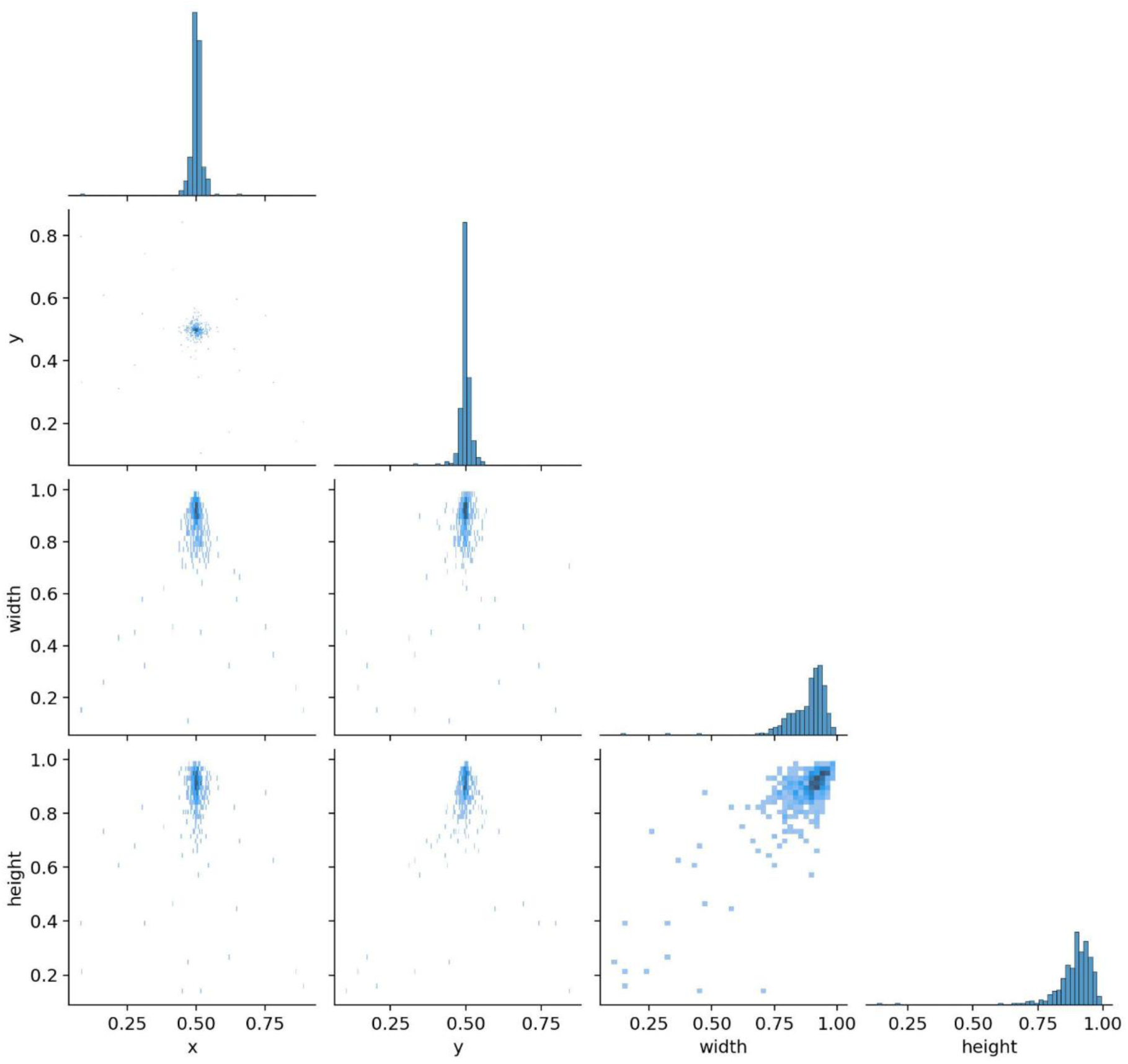
Distribution and Relationships of Bounding Box Parameters. Illustration of the distribution and pairwise relationships of bounding box parameters (x, y, width, height) generated by the YOLOv11m-seg model during lung tissue segmentation. The diagonal panels display histograms representing the marginal distributions of each parameter. The off-diagonal panels show scatter plots illustrating the pairwise relationships between each combination of parameters, with point density indicating frequency. The x and y parameters represent the normalized center coordinates of the bounding boxes, while width and height represent the normalized dimensions. The histograms reveal a central tendency for x and y coordinates around 0.5 and a bias towards larger width and height values. The scatter plots demonstrate the correlations between these parameters, providing insights into the spatial and dimensional characteristics of the detected lung tissue regions.

**Figure S10.**
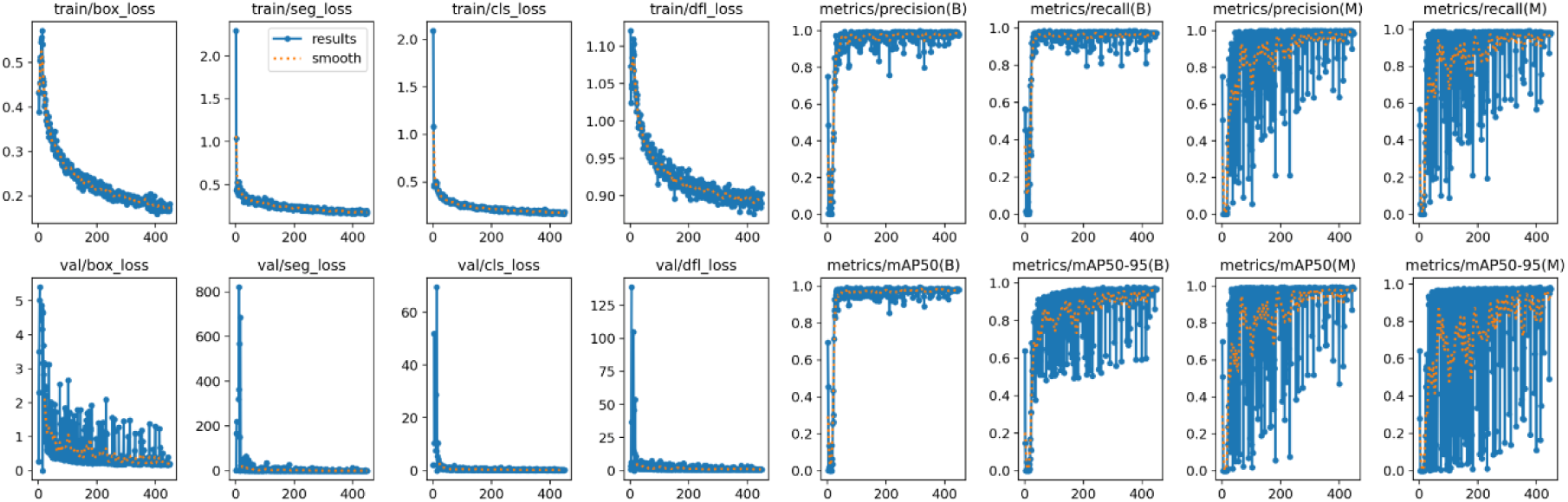
Training Dynamics of the YOLOv11m-seg Model. A comprehensive visualization of the model’s learning process, showcasing both the reduction in errors and the improvement in performance over time as shown in the comparison between training and validation metrics. The top row of the figure tracks the training losses for train/box_loss, train/seg_loss, train/cls_loss, and train/dfl_loss, where decreasing values indicate improved model learning. Simultaneously, the top row also displays the metrics/precision(B), metrics/recall(B), metrics/precision(M), and metrics/recall(M) metrics, representing precision and recall for both bounding box (B) and mask (M) predictions, with higher values signifying greater accuracy. The bottom row mirrors the top row, showing the corresponding validation losses for val/box_loss, val/seg_loss, val/cls_loss, and val/dfl_loss, and mean Average Precision (mAP) values for metrics/mAP50(B), metrics/mAP50-95(B), metrics/mAP50(M), and metrics/mAP50-95(M). In each plot, the raw results are represented by a blue line labeled results, while a yellow dotted line labeled smooth illustrates a smoothed trend, assisting in the visualization of the overall learning trajectory. The x-axis of each plot represents the training epochs, and the y-axis indicates the corresponding metric or loss value.

**Figure S11.**
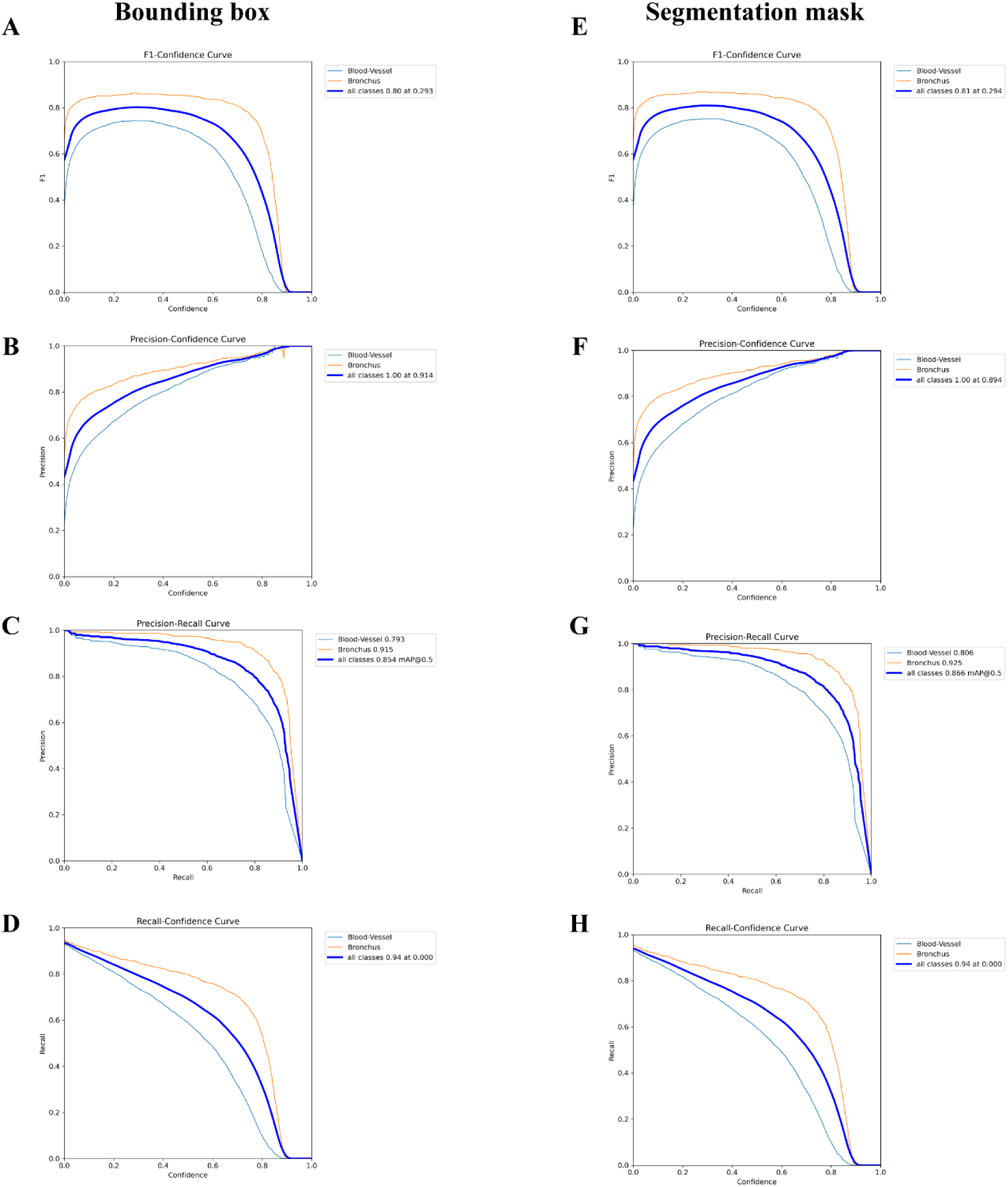
Performance Analysis of YOLOv11 Blood Vessel and Bronchus Segmentation. **(A-H)** The figure visualizes the YOLOv11 model’s performance in delineating blood vessels and bronchi, demonstrating its ability to maintain accuracy. **(A-D)** Illustrate bounding box prediction metrics, while **(E-H)** detail segmentation mask performance. The F1-Confidence curves **(A, E)** highlight the model’s balanced precision and recall, peaking at high F1 scores, indicating robust performance even at stringent confidence thresholds. The Precision-Confidence curves **(B, F)** show that the model consistently achieves high precision, reaching high precision at maximum confidence, signifying minimal false positives. The Precision-Recall curves **(C, G)** further confirm the model’s accuracy, demonstrating high precision values, illustrating its reliability in real-world applications. The Recall-Confidence curves **(D, H)** demonstrate the model’s sensitivity, maintaining high recall even at high confidence, ensuring that few relevant structures are missed.

**Figure S12.**
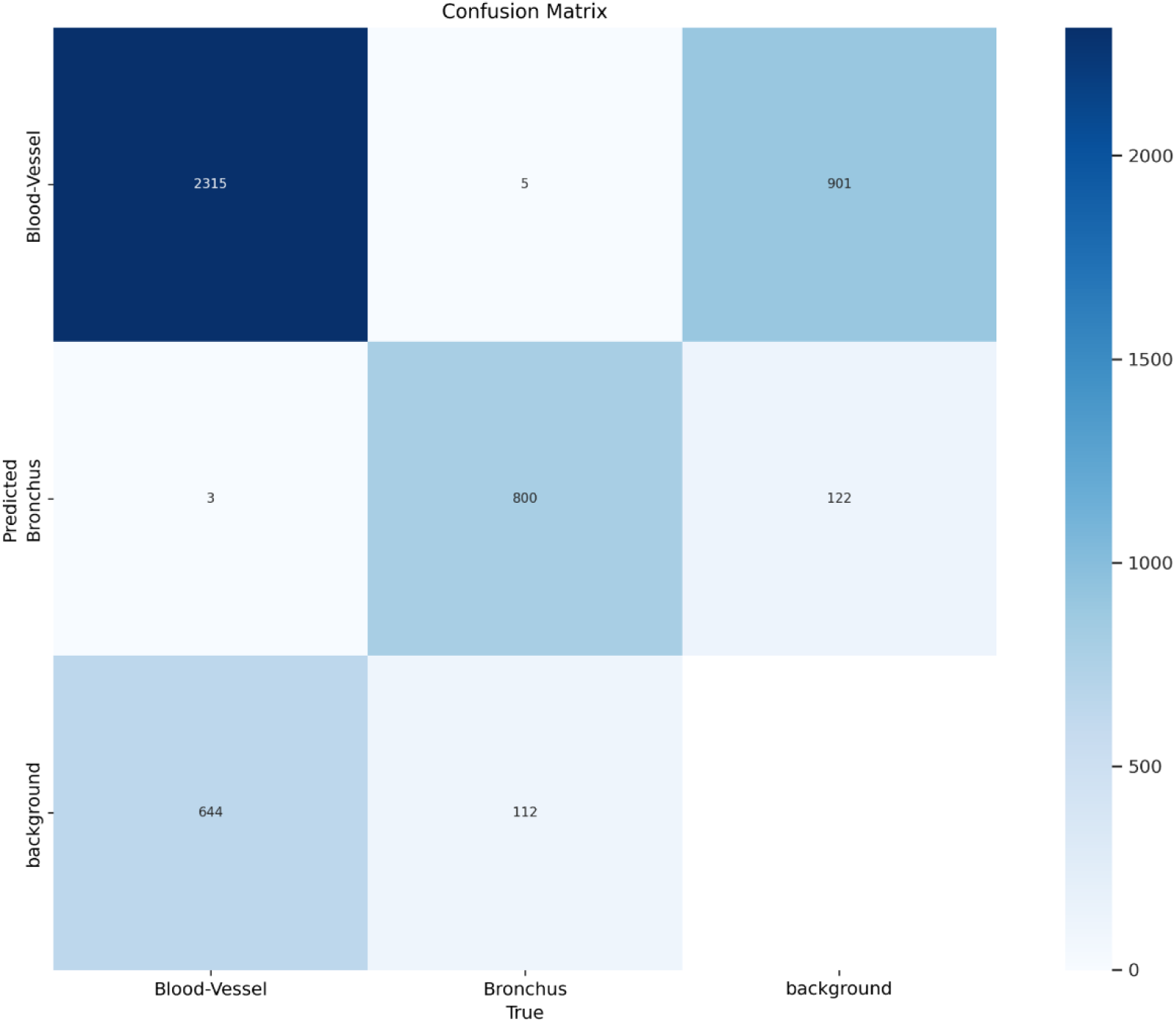
Segmentation Accuracy of Blood Vessel and Bronchus. Confusion matrix illustrates how well the model distinguished between blood vessels, bronchi, and the background during segmentation. It shows where the model’s predictions matched the actual labels and where it made mistakes. Each cell in the grid represents a category, with the rows showing what the model predicted and the columns showing what was present in the image. The numbers inside each cell indicate how many pixels fell into that category. For example, the model correctly identified 2315 pixels as blood vessels but misclassified 901 background pixels as blood vessels. Similarly, it correctly identified 800 pixels as bronchi but misclassified 122 background pixels as bronchi. The color scale on the right helps visualize the pixel counts, with darker colors indicating a higher number of pixels.

**Figure S13.**
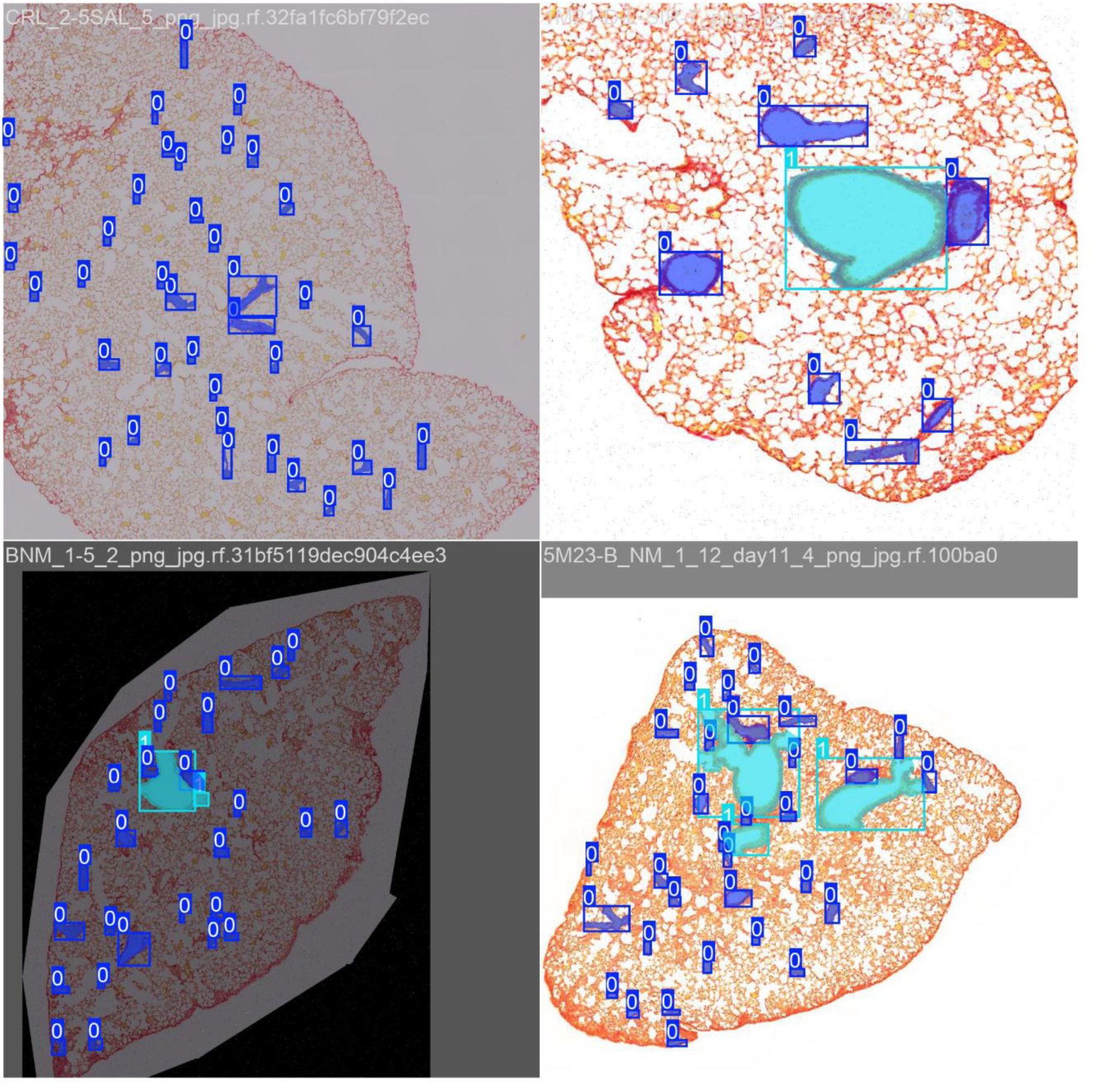
Visual Output of YOLOv11 Blood Vessel and Bronchus Detection During Training. Snapshot of the YOLOv11 model’s performance during the training phase, illustrating its ability to detect and segment blood vessels and bronchi within lung tissue histological images. Each panel displays a different image from the training batch, showcasing the model’s evolving accuracy. The images are overlaid with blue bounding boxes, which indicate the model’s detected regions of interest, highlighting where the model predicts the presence of either blood vessels or bronchi. Additionally, cyan segmentation masks represent the pixel-level segmentation of the detected structures, delineating the precise boundaries of blood vessels and bronchi as identified by the model. Each detected object is accompanied by a numerical label, representing the class identifier (e.g., 0 for blood vessels, 1 for bronchi). This visual representation offers insight into the model’s learning process, demonstrating its ability to progressively refine its detection and segmentation capabilities as it is trained on the histological image dataset. The figure highlights the model’s ability to identify and delineate these crucial lung tissue structures, which is essential for subsequent quantitative analysis.

**Figure S14.**
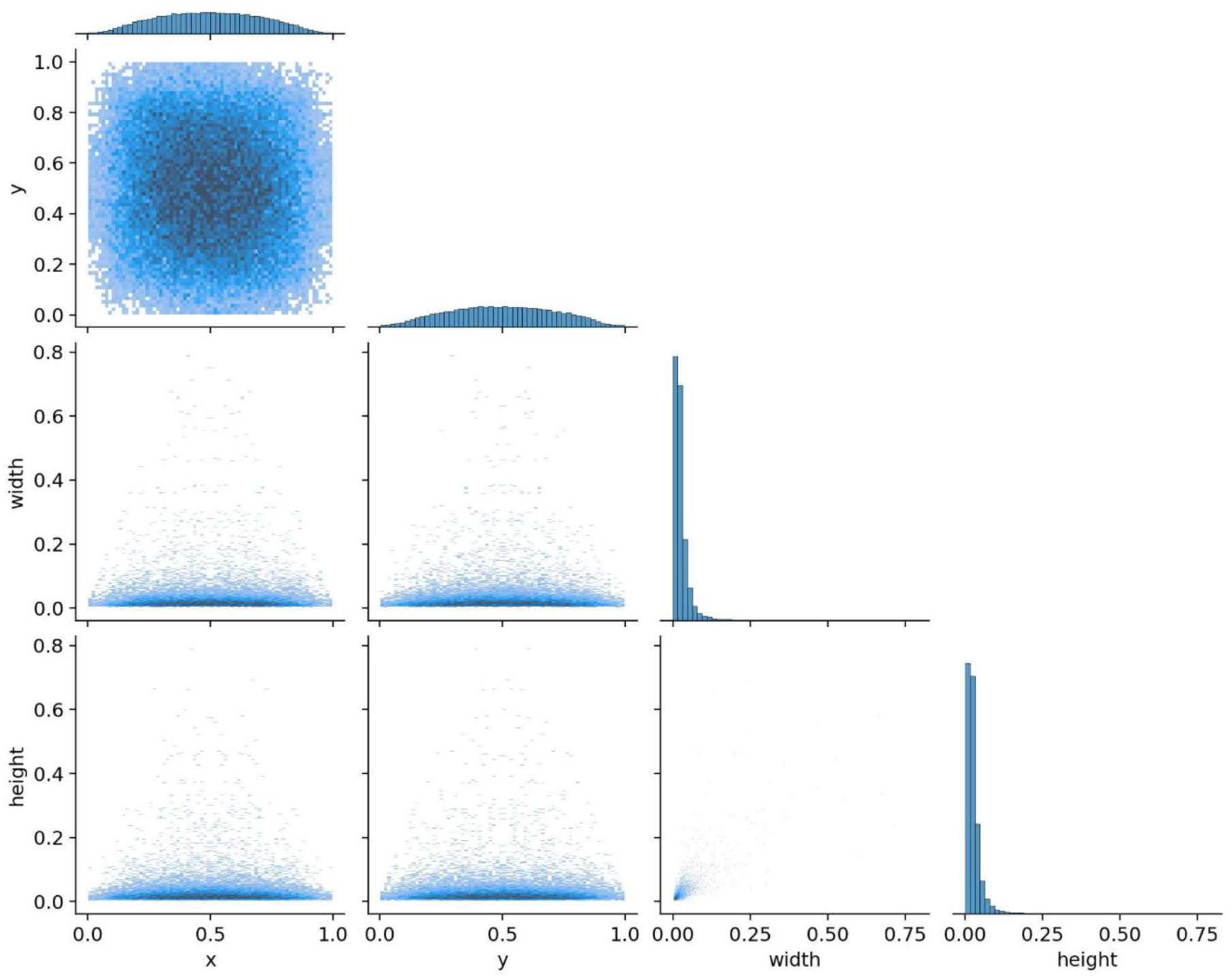
Distribution of Bounding Box Parameters for Blood Vessel and Bronchus Segmentation. An illustration of the bounding box parameters distribution resulting from the segmentation of blood vessels and bronchi. It displays histograms and scatter plots for the normalized horizontal centroid (x), normalized vertical centroid (y), normalized bounding box width, and normalized bounding box height. The diagonal panels show the frequency distribution of each individual parameter. The off-diagonal panels show scatter plots of pairs of these parameters.

**Figure S15.**
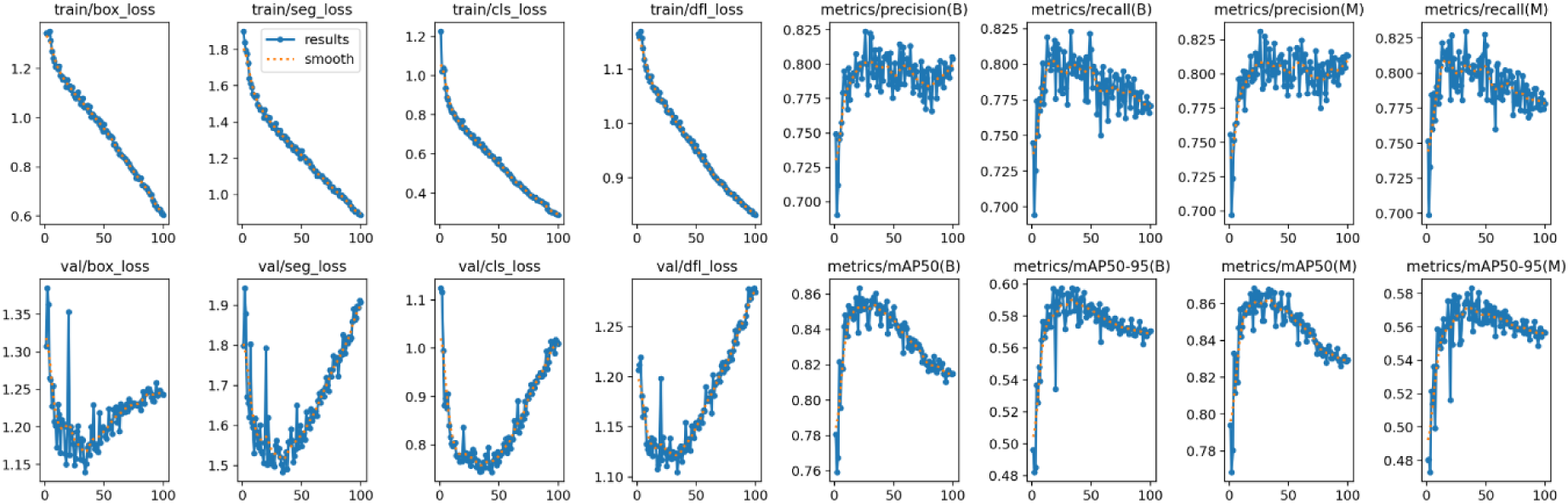
Model Training Performance and Loss Curves. Training and validation performance of the model during training over epochs. The top row shows training losses (box, segmentation, cls, and dfl) and training metrics (precision and recall for bounding boxes and masks). The bottom row displays corresponding validation losses and metrics (mAP50 and mAP50-95 for bounding boxes and masks). The dotted line in the training loss plots represents a smoothed version of the training loss.

**Table S5.** Performance and Operational Parameters of the YOLOv11m-seg Model. Comprehensive overview of the YOLOv11m-seg model’s performance and operational parameters. Module Information details the framework (Ultralytics 8.3.98), hardware (NVIDIA A100-SXM4-40GB), and software environment (Python 3.11.11, PyTorch 2.6.0+cu124, CUDA device 0) used for model execution. Performance metrics include box and mask precision (P), recall (R), and mean average precision (mAP 50-95), evaluating the model’s accuracy in object detection and segmentation. Speed parameters quantify computational efficiency through preprocessing, inference, postprocessing, and loss calculation times. Training Details summarize training parameters, including epochs, duration, learning rates, batch size, image size, and automatic mixed precision. Results Location specifies the saved model weights (best.pt, last.pt).

| Category | Metric | Value |
| --- | --- | --- |
| Modle Information | Model | YOLOv11m-seg |
|  | Framework | Ultralytics 8.3.98 |
|  | Neural network layers | 138 |
|  | Neural network parameters | 22,336,083 |
|  | Gradients | 0 |
|  | Giga Floating Point Operations per Second (GFLOPs) | 123 |
|  | Hardware | NVIDIA A100-SXM4-40GB (40,507 MiB) |
|  | Python Version | 3.11.11 |
|  | PyTorch Version | 2.6.0+cu124 |
|  | CUDA Device | 0 |
| Performance | Box Precision (P) | 0.986 (98.6%) |
|  | Box Recall (R) | 0.978 (97.8%) |
|  | Box mAP<sub>(50-95)</sub> | 0.978 (97.8%) |
|  | Mask Precision (P) | 0.993 (99.3%) |
|  | Mask Recall (R) | 0.985 (98.5%) |
|  | Mask mAP<sub>(50-95)</sub> | 0.98 (98%) |
| Speed | Preprocessing Time | 0.1 ms |
|  | Inference Time | 2.2 ms |
|  | Loss Calculation Time | 0.0 ms |
|  | Postprocessing Time | 0.8 ms |
| Training Details | Epochs Completed | 449/1000 |
|  | Total Training Time | 1.117 hours |
|  | Patience | 100 |
|  | Batch | 64 |
|  | Initial learning rate (lr0) | (1x10)<sup>(-4)</sup> |
|  | Final learning rate (lrf) | (1x10)<sup>(-3)</sup> |
|  | Image size (imgsz) | 640 |
|  | Automatic Mixed Precision (amp) | True |
| Results Location | Saved Weights | best.pt |
|  |  | last.pt |

**Table S6.** Lung Segmentation Validation, YOLOv11m-seg Performance Metrics. Validation results of the YOLOv11m-seg model for lung tissue segmentation, evaluated on 145 images containing 148 lung instances. Metrics include precision (P), recall (R), and mean average precision (mAP), reported for both bounding box and segmentation mask predictions. The model achieved box P=0.986, R=0.978, mAP (50–95) =0.978, and mask P=0.993, R=0.985, mAP50=0.993, mAP (50–95)=0.98, demonstrating high accuracy in lung tissue segmentation.

| Class Name | Images | Instances | Box<br>Precision (P) | Box<br>Recall (R) | Box<br>mAP<sub>(50-95)</sub> | Mask<br>Precision (P) | Mask<br>Recall (R) | Mask<br>mAP<sub>50</sub> | Mask<br>mAP<sub>(50-95)</sub> |
| --- | --- | --- | --- | --- | --- | --- | --- | --- | --- |
| Lung | 145 | 148 | 0.986 | 0.978 | 0.978 | 0.993 | 0.985 | 0.993 | 0.98 |

**Table S7.**
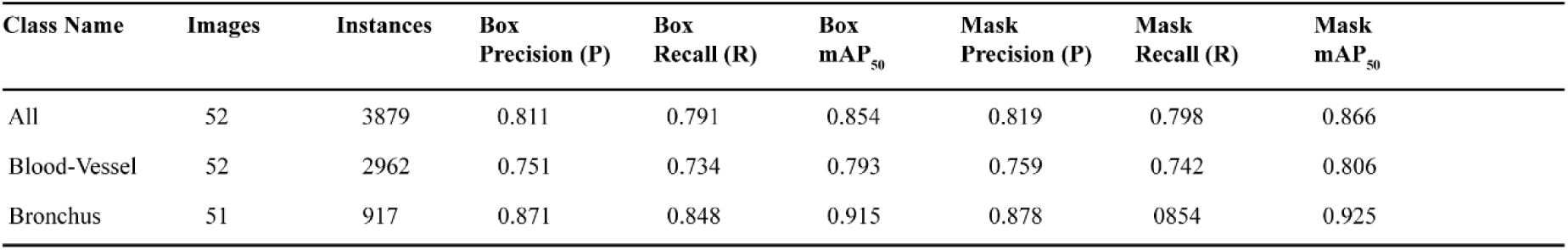
Blood Vessel and Bronchus Segmentation Validation, YOLOv11 Performance Metrics. Table show the validation outcomes of the YOLOv11 model applied to blood vessel and bronchus segmentation within lung tissue histological images. The model was trained on 260 images, but due to computational constraints, the validation was performed on batches of 50 images, with the final reported metrics based on a subset of 52 images. Performance was assessed across three categories: all detected objects, specifically blood vessels, and specifically bronchi. Metrics were calculated for both bounding box and segmentation mask predictions, including precision, recall, and mean average precision at 0.5 threshold. The overall performance, considering all classes, demonstrated bounding box metrics of 0.811 precision, 0.791 recall, 0.854 mAP50, with corresponding mask metrics of 0.819 precision, 0.798 recall, 0.866 mAP50. For blood vessel segmentation, the model achieved bounding box metrics of 0.751 precision, 0.734 recall, 0.793 mAP50, and mask metrics of 0.759 precision, 0.742 recall, 0.806 mAP50. Bronchus segmentation yielded bounding box metrics of 0.871 precision, 0.848 recall, 0.915 mAP50, and mask metrics of 0.878 precision, 0.854 recall, 0.925 mAP50. These results, derived from the 52-image subset, illustrate the model’s capacity for accurate detection and segmentation of vascular and bronchial structures within the analyzed histological images, despite the validation being performed on a subset of the original dataset.

| Class Name | Images | Instances | Box Precision (P) | Box Recall (R) | Box mAP<sub>50</sub> | Mask Precision (P) | Mask Recall (R) | Mask mAP<sub>50</sub> |
| --- | --- | --- | --- | --- | --- | --- | --- | --- |
| All | 52 | 3879 | 0.811 | 0.791 | 0.854 | 0.819 | 0.798 | 0.866 |
| Blood-Vessel | 52 | 2962 | 0.751 | 0.734 | 0.793 | 0.759 | 0.742 | 0.806 |
| Bronchus | 51 | 917 | 0.871 | 0.848 | 0.915 | 0.878 | 0.854 | 0.925 |

**Table S8.** YOLOv11x-seg Model Summary and Performance Metrics of the blood vessels and bronchus segmentation. The table summarizes of the YOLOv11x-seg model. It shows the Model Information, such as the model’s architecture, framework, number of layers and parameters, hardware, and software versions. It also presents Performance metrics, like precision, recall, and mAP for bounding boxes and masks. The table includes Speed measurements, showing the time for preprocessing, inference, loss calculation, and postprocessing. Training Details are provided, including epochs, training time, patience, batch size, learning rates, and image size. Finally, the Results Location indicates where the model’s weights are saved.

| Category | Metric | Value |
| --- | --- | --- |
| Modle Information | Model | YOLOv11x-seg |
|  | Framework | Ultralytics 8.3.98 |
|  | Neural network layers | 203 |
|  | Neural network parameters | 62,004,438 |
|  | Gradients | 0 |
|  | Giga Floating Point Operations per Second (GFLOPs) | 318.5 |
|  | Hardware | NVIDIA H200 (141,132 MiB) |
|  | Python Version | 3.11.11 |
|  | PyTorch Version | 2.8.0+cu128 |
|  | CUDA Device | 0 |
| Performance | Box Precision (P) | 0.811 (81.1%) |
|  | Box Recall (R) | 0.791 (79.1%) |
|  | Box mAP<sub>(50-95)</sub> | 0.596 (59.6%) |
|  | Mask Precision (P) | 0.819 (81.9%) |
|  | Mask Recall (R) | 0.798 (79.8%) |
|  | Mask mAP<sub>(50-95)</sub> | 0.583 (58.3%) |
| Speed | Preprocessing Time | 0.3 ms |
|  | Inference Time | 30.4 ms |
|  | Loss Calculation Time | 0.0 ms |
|  | Postprocessing Time | 7.1 ms |
| Training Details | Epochs Completed | 100/100 |
|  | Total Training Time | 3.570 hours |
|  | Patience | 100 |
|  | Batch | 4 |
| | Initial learning rate (lr0) | $(1 \times 10)^{-4}$ |
| | Final learning rate (lrf) | $(1 \times 10)^{-2}$ |
|  | Image size (imgsz) | 3840 |
|  | Automatic Mixed Precision (amp) | True |
| Results Location | Saved Weights | best.pt<br>last.pt |

**Table S9.**
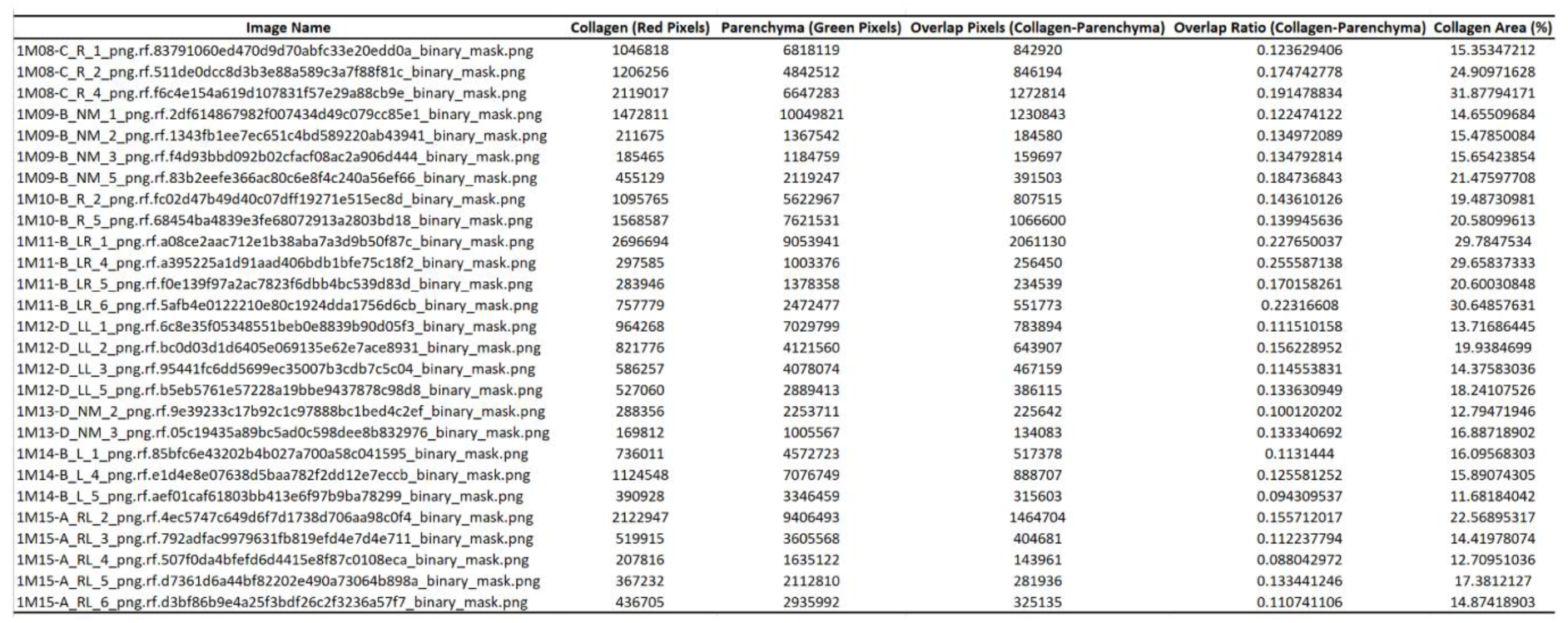
Quantitative Analysis of Collagen and Parenchyma Distribution. Quantitative analysis of collagen and parenchyma distribution within tissue samples. The table provides several key parameters: Image Name identifies each analyzed image, Collagen (Red Pixels) quantifies the number of pixels identified as collagen, and Parenchyma (Green Pixels) quantifies the number of pixels identified as parenchyma. Overlap Pixels (Collagen-Parenchyma) indicates the number of pixels where collagen and parenchyma masks overlap, representing their spatial co-localization. Overlap Ratio (Collagen-Parenchyma) expresses the percentage of collagen area overlapping with parenchyma area, and Collagen Area (%) represents the percentage of the total tissue area occupied by collagen.

## Supplementary Methods

### Fixation and paraffin embedding

Tissue samples were fixed in 4% PFA and washed three times with PBS. Following fixation, samples underwent routine dehydration and clearing using a tissue processor (Leica TP1020, Germany).

Processed tissues were subsequently embedded in Paraplast paraffin (Leica) using the Histocore Arcadia H embedding system (Leica) to generate solid paraffin blocks.

Paraffin blocks were sectioned at 4μm thickness using a rotary microtome. Sections were floated on a 37-39°C water bath for optimal flattening and mounted onto charged glass slides. Slides were dried overnight at 37 °C.

### Picrosirius Red staining protocol for lung paraffin sections

Paraffin-embedded lung sections were stained with Picrosirius Red to visualize collagen deposition. Slides were first deparaffinized in xylene for 5 min, three times, and then rehydrated through a graded ethanol series consisting of 100% ethanol for 1 min, three times, followed by 95% ethanol for 1 min and 80% ethanol for 1 min. Slides were then rinsed in running tap water for 1 min.

Sections were stained with Weigert’s iron hematoxylin for 9 min, followed by washing in running tap water for 5 min, or until excess stain was completely removed and the sections appeared clear. Collagen staining was performed by incubating the slides in Picrosirius Red solution for 1 h. Excess dye was removed by rapid differentiation in 0.5% acetic acid solution, using three quick dips repeated twice. Slides were then rapidly dehydrated in 100% ethanol using three quick dips repeated twice, cleared in xylene for 1 min three times, and held in the final xylene wash until mounting.

Weigert’s iron hematoxylin was prepared by mixing Solution A and Solution B at a 1:1 ratio immediately before use. Solution A was prepared by dissolving 5 g hematoxylin in 500 mL of 95% ethanol and can be stored for up to 1 year. Solution B was prepared by mixing 20 mL of 29% aqueous ferric chloride solution, 475 mL double-distilled water, and 5 mL of 32% hydrochloric acid, and can be stored for up to 1 year. The 29% aqueous ferric chloride solution was prepared by dissolving 29 g ferric chloride in 100 mL double-distilled water and can be stored for up to 1 year.

To prepare 260 mL of working Weigert’s iron hematoxylin solution, 130 mL of Solution A was mixed with 130 mL of Solution B. The working solution can be used for up to 4 days. Picrosirius Red solution was prepared by dissolving 0.25 g Direct Red 80 in 250 mL picric acid solution, followed by thorough mixing and filtration. The solution can be stored for up to 3 years. A 0.5% acetic acid solution was prepared by adding 5 mL glacial acetic acid to double-distilled water and bringing the final volume to 1000 mL.

## Notes

https://github.com/Anas-Odeh/FibroSight

### Competing Interest Statement

The authors have declared no competing interest.

